# GiPS: Genomics-informed parent selection uncovers the breeding value of wheat genetic resources

**DOI:** 10.1101/2021.12.15.472759

**Authors:** Albert W. Schulthess, Sandip M. Kale, Fang Liu, Yusheng Zhao, Norman Philipp, Maximilian Rembe, Yong Jiang, Ulrike Beukert, Albrecht Serfling, Axel Himmelbach, Jörg Fuchs, Markus Oppermann, Stephan Weise, Philipp H. G. Boeven, Johannes Schacht, C. Friedrich H. Longin, Sonja Kollers, Nina Pfeiffer, Viktor Korzun, Matthias Lange, Uwe Scholz, Nils Stein, Martin Mascher, Jochen C. Reif

**Affiliations:** Leibniz Institute of Plant Genetics and Crop Plant Research (IPK) Gatersleben, Seeland, Germany; Carlsberg Research Laboratory, Copenhagen, Denmark; Julius Kühn Institute (Federal Research Centre for Cultivated Plants), Quedlinburg, Germany; Limagrain GmbH, Peine-Rosenthal, Germany; State Plant Breeding Institute, University of Hohenheim, Stuttgart, Germany; KWS SAAT SE & Co. KGaA, Einbeck, Germany; KWS LOCHOW GmbH, Bergen, Germany; Center for Integrated Breeding Research (CiBreed), Georg-August-University, Göttingen, Germany; German Centre for Integrative Biodiversity Research (iDiv) Halle-Jena-Leipzig, Leipzig, Germany

## Abstract

The great efforts spent in the maintenance of past diversity in genebanks are rationalized by the potential role of plant genetic resources in future crop improvement – a concept whose practical implementation has fallen short of expectations. Here, we implement genomics-informed parent selection to expedite pre-breeding without discriminating against non-adapted germplasm. We collect dense genetic profiles for a large winter wheat collection and evaluate grain yield and resistance to yellow rust in representative coresets. Genomic prediction within and across genebanks identified the best parents for PGR x elite derived crosses that outyielded current elite cultivars in multiple field trials.

## Main

Genebanks around the world are committed to maintain permanently plant genetic resources (PGR), some of which have not been grown on farmer’s field for a century or, in the case of crop-wild relatives, have never been used as crops at all^1^. PGR underperform dramatically in current agricultural environments^2^. Most of them, for example, succumb to pathogens currently at large, preventing an un-biased assessment of their breeding value^3^. Pre-breeders have often ended up in choosing “exotic” genotypes too closely related to the elite genepool or with the inadvertent loss of novel haplotypes by selection in the field^4,5^. We and others^6,7^ have bemoaned the disconnect between genebank management and breeding resulting from a lack of effective and generally applicable strategies to identify valuable germplasm as donors in pre-breeding programs.

In recent years, genomic approaches have showcased the potential of exotic germplasm for plant breeding^4,5,8^. Pan-genomes^9^, rapid gene cloning and targeted enrichment sequencing^10^ have accelerated the isolation of resistance genes, including those from crop-wild relatives. However, their durable deployment requires either complex pyramiding schemes or transgenic methods^11^ under tight regulation in many countries. Moreover, resistance gene breeding takes advantage of a comparatively simple genetic architecture where single, easily transferable major genes confer large genetic gains^12^. After the Green Revolution^13^, reductionist approaches to further increase grain yield per se, a genetically complex trait with high genotype-by-environment interaction, have been counter-productive in practice^14,15^.

A combination of genebank genomics (genome-wide marker profiles for entire genebank collections) and genomic prediction (inference of phenotype from genotype) has been proposed as one way forward to characterize and prioritize genebank accessions for pre-breeding^16^. The missing link is the accurate phenotyping of a training set from which the breeding value of thousands of accessions can be predicted. We have proposed a hybrid strategy, in which agronomic performance is not scored in the PGR itself (“*per se*”), but in a hybrid ‘Elite x PGR’ background to negate the masking effect of a lack of agronomic adaption of germplasm preserved ex *situ* or locally abandoned^3^. However, the high cost of large-scale cross-fertilization in inbreeding crops^17^ and the low heritability estimated in a hybrid context, left reasonable doubts as to whether phenotypic evaluation in a panel, small enough to be tractable for hybrid production, can inform genomic prediction of breeding in thousands of accessions. Here, we report on the implementation and evaluation of our strategy in winter wheat, the most important food crop in Europe.

## Results

### Genetic structure of a global winter wheat collection

The universe of genetic and phenotypic diversity from which we selected parents for pre-breeding traces back to 7,651 wheat accessions from the German Federal ex *situ* Genebank (**Supplementary Table 1**). Genotypic characterization was performed on genotypes descended from single spikes to reduce the effect of heterogeneity within accessions. Genotyping-by-sequencing (GBS) of this panel (henceforth the “IPK collection”) together with 325 European elite cultivars yielded 69,356 bi-allelic single-nucleotide polymorphism (SNP) markers with less than 10% missing genotype calls (**Supplementary Table 2**). A very small fraction of accessions (0.2 %) turned out to be mis-classified tetraploid wheats (confirmed by flow cytometric ploidy analysis and the most recent passport records), illustrating the value of genome-wide marker profiles for the curation of genebank collections^8,18,19^ (**Supplementary Tables 3** and **4** and **Supplementary Figs. 1** and **2**). Consistent with previous large-scale genebank genomics efforts^18,19^, duplicates abound in the IPK collection: 37 % of accessions were highly similar (> 99 %) to at least one other accession (**Supplementary Table 5**).

We intersected our GBS-based variant calls with high-density SNP genotyping data of 2,608 winter wheat accessions from the French national genebank^20^ (the “INRAE collection”), yielding 895 shared variants. A high correlation of distance measures (r = 0.82, Mantel p-value < 0.01) indicates that rather few common markers can still accurately capture population structure (**Supplementary Table 6**). Seventeen percent of INRAE accessions had a close (> 99 % identity) match to at least one IPK accession (**Supplementary Table 7**). By contrast, 434 INRAE accessions did not match to any IPK accessions (**Supplementary Table 8**), while vice versa, 2,161 IPK accessions lacked counterparts in the INRAE collection (**Supplementary Table 9**). These observations reinforce calls to action demanding across-genebank curation efforts informed by genomics (https://agent-project.eu/).

A joint principal component analysis (PCA) revealed that both genebanks largely covered the same diversity space (**Fig. 1a**). A PCA highlighted two major germplasm groups, corresponding to accessions of European and Asian origin, respectively (**Fig. 1b,c**). Southern European accessions occupied an intermediate position. Interestingly, wheats from Western Europe and Eastern Europe were well separated in the diversity space. Most accessions from the Americas were descended from the Eastern European genepool. Genetic differentiation as measured by F_ST_ was consistent with the patterns evident from the PCA (**Fig. 1d**). The pronounced genetic divergence (F_ST_ = 0.376) between European elite cultivars and the more diverse Asian germplasm (**Fig. 1e**) makes it likely that the targeted use of Asian accessions in European wheat breeding may reap the same rewards as an exchange of alleles in the opposite direction^21^.

**Fig. 1.**
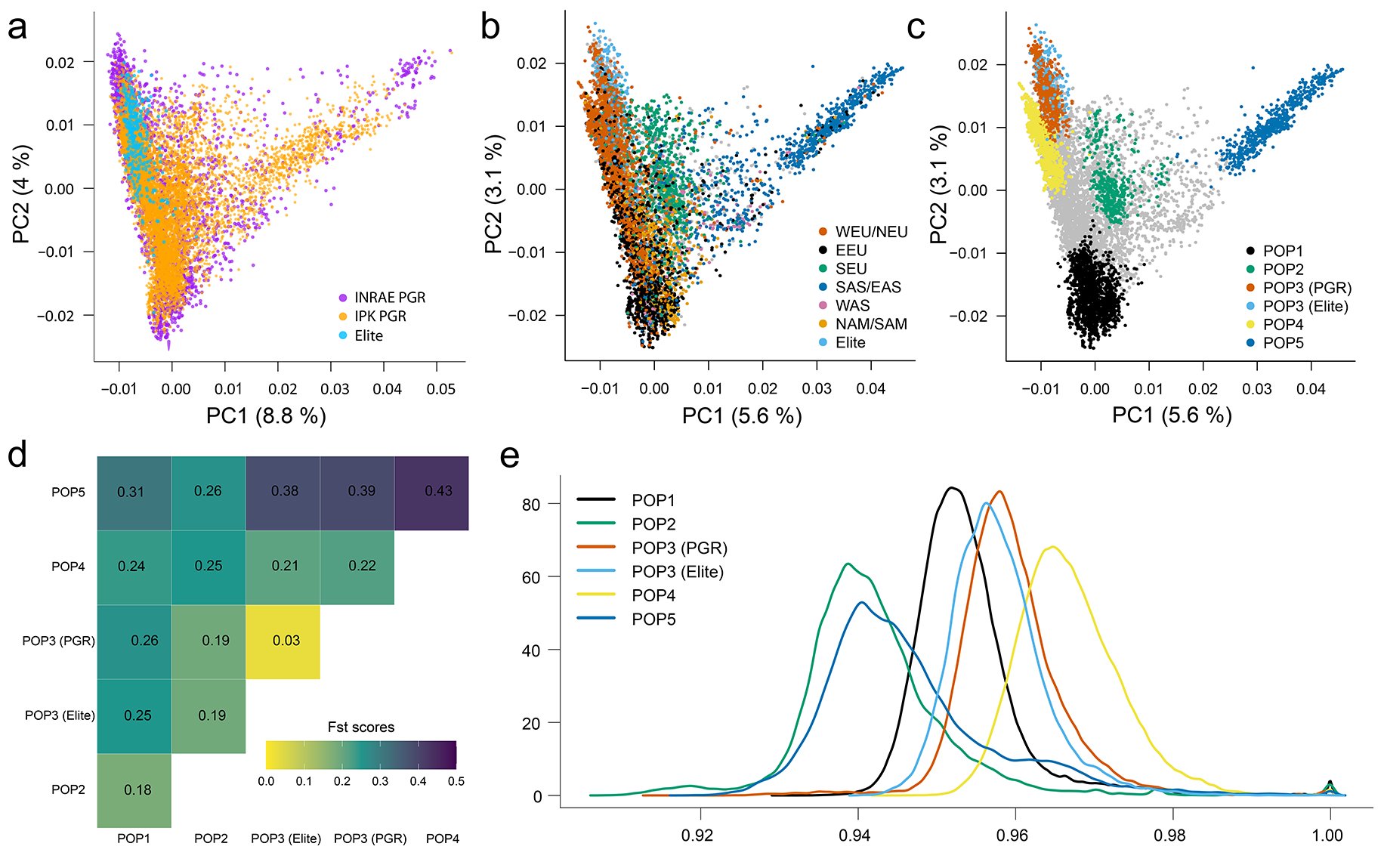
Genetic diversity within and between genebanks. **a**, Principal component analysis (PCA) representing the diversity space of 2,608 winter wheat accessions from INRAE (blue color) onto which 7,745 IPK genebank samples (red color) and 325 European elite cultivars (Elite) were projected. **b**, PCA plot depicting genetic diversity harvored by IPK genebank accessions and elite cultivars. The IPK accessions are colored based on different geographic regions: WEU/NEU, western/northern Europe; EEU, Eastern Europe; SEU, Southern Europe; SAS/EAS, Southern/Eastern Asia; WAS, Western Asia; NAM/SAM, North/South America. Samples from underrepresented regions are presented in gray. **c**, The same data as in **(b)** are shown, but samples are colored by their assignments to subpopulations defined by ADMIXTURE (k=5). Samples with ancestry coefficients <0.7 are shown in grey. Subpopulations 1-5 were mostly conformed (> 50% known origins) by genotypes from EEU, SEU, WEU/NEU, WEU/NEU (mostly Germany) and SAS/EAS, respectively. **d**, Fixation index (F_st_) between subpopulations defined by ADMIXTURE. **e**, Distributions of similarity within subpopulations. The density of pairwise identity-by-state values is plotted.

### Definition of trait-customized core collections

After a thorough inspection of patterns of diversity, we aimed at singling out genotypes that would stand a high chance of increasing crop performance when crossed with current elite varieties. The first trait we focused on is resistance to yellow rust (YR) caused by *Puccinia striiformis* f. sp. *tritici* (**Supplementary Tables 10**-**12**). As expected, the vast majority of genebank accessions (91.6 %) was more susceptible to naturally occurring YR than elite cultivars (**Supplementary Figs. 3** and **4**). Solely focusing on highly resistant accessions incurs the risk of enriching for alleles from the modern European genepool already deployed by breeders to the detriment of resistance-conferring alleles contributed by non-adapted landraces from other genepools (**Supplementary Tables 13** and **14**). Based on preliminary data, we compiled a “trait-customized” core collection (T3C) of diverse accessions with 150 genotypes resistant to YR and 50 closely related susceptible genotypes to strike a balance between power in association mapping, allelic richness and coverage of the diversity space (see **Supplementary Note** and **Supplementary Fig. 5**). The same strategy was applied to address the problem for two other important diseases: leaf rust and powdery mildew. The three T3Cs together with a coreset of 200 diverse elite cultivars were intensively tested relying on natural and artificial YR infections in multiple field trials, which provided a very high trait heritability (**Supplementary Tables 15**-**17**). Whole genome re-sequencing data (3-fold coverage) was generated for 263 genotypes of the three T3Cs and 191 elite cultivars (**Supplementary Table 15**). Read mapping and SNP calling against the Chinese Spring Reference^22^ provided a dataset of 2,788,918 SNPs with less than 10% missing calls, which was then used to perform genome-wide association scan (GWAS) analysis and genomics-informed parent selection (GiPS) for resistance breeding (see results after the next section).

### Atlas of footprints of selection in the European elite pool

Genome scans for regions under selection can reveal loci targeted by breeders as well as wider regions which are in linkage to selected loci with reduced haplotype diversity^23^. To find genomic footprints of selection in European winter wheat, we expanded our whole genome shotgun data with diversity from 183 additional accessions and 131 modern German breeding lines. According to their acquisition/release years, a total 760 genotypes constitute roughly a historic time course (**Supplementary Table 18**): 255 genotypes predating the Green Revolution (PreGreen), 212 varieties released between 1971 and 2000 (OldCV), and 293 recent genotypes bred after 2000 (NewCV). A scan for high cross population composite likelihood ratios (XP-CLR)^24^ revealed 1,304, 1,201, and 1,001 selective sweep regions between PreGreen-and-OldCV, OldCV-and-NewCV, and PreGreen-and-NewCV combinations, respectively (**Supplementary Table 19**). Several selective sweep regions (XP-CLR score ≥ 40) were co-located with known disease resistance loci (**Fig. 2a**) such as *Lr10* on chromosome 1AS^25^ and *Yr17* on chromosome 2AS^26,27^. The latter, introgressed from *Aegilops ventricosa*^9^, illustrates how alien introgressions have been successfully used in wheat breeding^28^ and stimulated a systematic scan for the presence of alien chromatin in our historic panel. To this end, we employed approaches based on *k*-mers and read depth (**Fig. 2b,c**, see **Online Methods**). We detected seven previously described^9,23^ introgressions on chromosomes 2A, 4A, 1B, 2B, 2D, 3D, and 7D (**Supplementary Table 20**). Moreover, we found evidence for hitherto unknown alien introgressions on chromosomes 1D, 2D and 5B. We compared the potentially novel introgressed haplotypes with published genomes of wheat wild relatives^9^, but were not able to unravel their origins. Elite cultivars being currently grown in Europe may harbor multiple independent introgressions (**Fig. 2d**), with up to six introgressions in modern cultivars such as ‘Anapolis’ and ‘Memory’ (**Supplementary Table 21**). Strikingly, the frequency of all introgressions has increased in recent decades (**Fig. 2e)**, with the notable exceptions of 1D and the 1BS-1RS whole-arm introgression dating back to the 1920s^29^. Despite conferring multiple disease resistances^30,31^ and abiotic stress tolerance^32^, the appeal of 1BS-1RS to breeders may have waned due to new pathotypes that overcame resistances^31,33^, or because of its negative effects on bread-making quality^34^. This clearly illustrates how the popularity of introgressions is tightly linked to their overall net value for breeders.

**Fig. 2.**
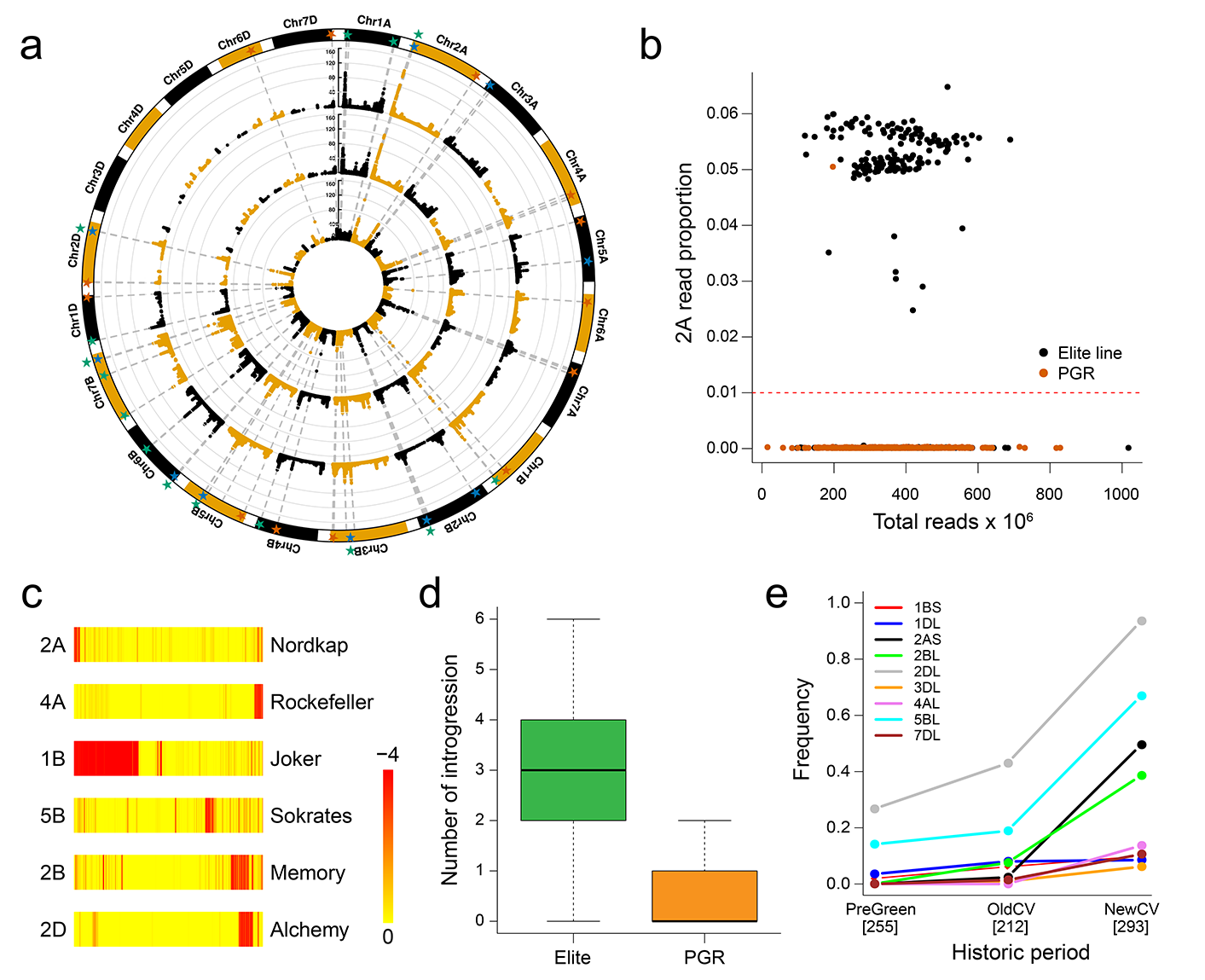
Tracing the history of introgression breeding. **a**, Circos plot of normalized XP-CLR scores for contrasts between three different historic periods: PreGreen (before 1970), OldCV (1971-2000) and NewCV (2001-Present). The regions under selection are highlighted with stars. Inner circle: PreGreen vs OldCV (red stars); middle circle: OldCV vs NewCV (blue stars); outer circle: PreGreen vs NewCV (green stars). **b**, Detection of introgressions by measuring the abundance of diagnostic *k*-mers. The relative frequency of *k*-mers specific to the *Ae. ventricosa* introgression on chromosome 2A are shown for plant genetic resources (PGR) and elite lines. The introgression is more frequent in the elite panel. **c**, Drops in read depth (log_2_-fold changes compared to Chinese Spring) are indicative of introgressions. Read depth in 100 kb along six chromosomes is shown for one putative introgression carrier. Most introgressions occur in distal regions. **d**, Number of putative introgressions detected in 318 diverse elite lines and 442 diverse PGR samples. **e**, Frequencies of selected introgressions according to the three different historic periods. The number of varieties per period are indicated in brackets [ ].

### Genome-wide association mapping for yellow rust

GWAS is used to detect markers and haplotypes linked to agronomic traits and, in the best case, can provide candidate gene resolution^35^. Pan-genomic infrastructures, however, may be needed to obtain the full gene complement content under GWAS peaks, in particular as families of the major classes of candidate resistance genes are subjected to abundant structural variation^36,37^. To deal with a possible overcorrection of associations due to genetic differentiation between elite cultivars and accessions (**Fig. 3a,b**), GWAS was conducted at three different levels: whole set, only elite cultivars and only T3C accessions. In this way, association scans for YR detected 684, 194, and 29 significantly [−log_10_(p-value) > 5.97] associated SNPs in the whole set, T3C accessions and elite cultivars population, respectively (**Fig. 3c-e** and **Supplementary Table 22**). Some associated regions were co-located with known, but not yet cloned resistance genes loci such as *Yr17* (2AS)^26,27^ and *Yr75* (7AL)^38^, which have been widely deployed in elite cultivars (**Fig. 3c-e** and **Supplementary Table 23**). We observed long haplotype blocks under GWAS peaks in elite cultivars, which can likely be attributed to intense and effective selection by breeders for major genes (**Supplementary Fig. 6**), and which complicate the prioritization of candidate genes. By contrast, linkage disequilibrium decays in general much faster around GWAS peaks detected in PGR (**Supplementary Table 24**), possibly increasing mapping resolution once pan-genome assemblies of resistant haplotypes become available.

**Fig. 3.**
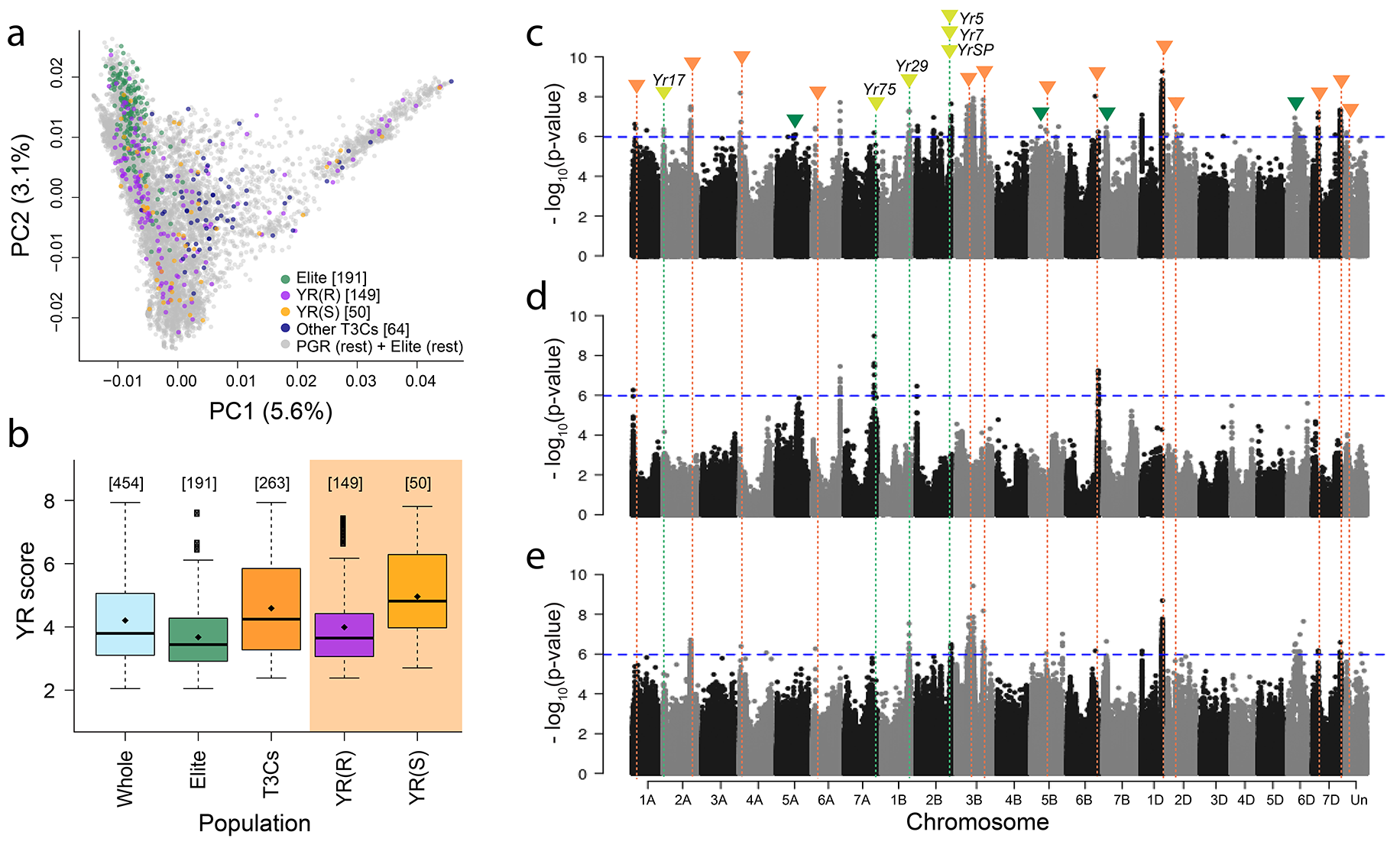
Deep mining plant genetic resources (PGR) of the IPK genebank for new sources of resistance against yellow rust (YR) not yet used in winter wheat breeding. **a**, Molecular diversity potrayed by the first two principal components (PC) from genotyping-by-sequencing variants and covered by the mined panel: 191 diverse elite cultivars, 199 diverse PGR [YR(R) plus 50YR(S)] from a YR trait-customized core collection (T3C) selected for resistant PGR enrichment and minimized population structure, plus 64 diverse PGR from T3Cs for other diseases. **b**, YR score (1 = fully resistant, 9 = fully susceptible) of the mined panels based on intensive field phenotyping. Diamonds indicate the average of distributions, while numbers in brackets [ ] in (**a**) and (**b**) indicate the number of datapoints for each category. Genomic regions harboring loci associated to YR resistance as revealed by whole genome sequencing and association mapping in (**c**) the whole panel (elites + T3Cs PGR), (**d**) only elites and (**e**) only T3Cs PGR. Physical positions are according to the Chinese Spring v1.0 reference. Significance thresholds are denoted with blue horizontal dashed lines. Positions of known YR resistance loci are indicated with lemon green vertical dashed lines and triangles, while green triangles denote hitherto not known resistance loci with resistance conferring alleles almost fixed in elites. Regions harboring new sources of resistance and fully contributed by PGR are shown with orange dashed lines and triangles.

To understand the contribution of structural variation to the genetic architecture of disease resistances, we conducted *k*-mer GWAS, which uses the presence-absence state of short sequence fragments of fixed length (*k*-mers) as a proxy for structural variants (see **Online Methods**). The most highly associated *k*-mers, when mapped to the Chinese Spring genome, were co-located with peak regions identified in SNP-GWAS (**Supplementary Fig. 7**). A notable exception was the 2AS (*Yr17*) peak, which was much less prominent in the SNP-GWAS, likely because of shortcomings of the reference-based SNP calling with highly diverse alien haplotypes. Importantly, 142 of 533 significantly associated *k*-mers were absent from the wheat pan-genome assemblies^9^ (**Supplementary Table 25**). Genome assemblies of accessions harboring these *k*-mers (**Supplementary Table 26**) and the resistant haplotypes they represent are priority targets for expanding the wheat YR resistance atlas^39^ and the global wheat pan-genome infrastructure.

GWAS peaks that do not correspond to known resistance genes may tag beneficial haplotypes from landraces that have not yet been deployed by breeders. However, resistance-conferring alleles for SNPs underlying associations on chromosomes 5A, 5B, 6D, 7B and one additional SNP on the unassigned chromosome were nearly fixed (allele frequency > 98%) in elite cultivars (**Fig. 3c-e** and **Supplementary Table 22**), suggesting that these putative resistance loci, despite their importance in breeding, have not been scrutinized in prior genetic studies. Resistance-conferring alleles private to PGR were found for 606 associated SNPs on chromosomes 1A, 1B, 1D, 2A, 2B, 2D, 3B, 4A, 5B, 6A, 6B and 7D, plus one SNP mapping to sequences not assigned to chromosomes (**Fig. 3c-e** and **Supplementary Table 22**). Taken together, these loci point to 30 resistance-conferring haplotypes absent from elite cultivars (**Supplementary Fig. 8** and **Supplementary Table 27**). Genotypes carrying such resistance-conferring haplotypes at a single locus, but otherwise having a susceptible genomic background are approximately equivalent to near-isogenic lines (NIL) and as such will be good starting points for genetic and functional characterization. A haplotype analysis for all GWAS peaks indicated that such NIL proxies are rare among PGR: on average, the 23 potential donors carry 24 from a total of 66 resistance-conferring haplotypes, while more than half the donors carry more than one potential source of novel resistances (**Supplementary Table 27**). By contrast, promising donors for breeding programs carrying many resistance-conferring alleles are easy to pick (**Supplementary Table 28**). A notable example is the Iranian landrace TRI 5804 bearing 28 of the 30 possible novel resistance-conferring alleles.

### Genomics-informed parent selection (GiPS)

The widespread use of alien introgressions has proven that beneficial genes and haplotypes from PGR can be identified and deployed effectively. The perseverance and good luck of pre-breeders in over-coming crossing barriers may have played a large role in the success of crop-wild relative introgressions in wheat breeding^23^. Can the same be achieved with (new)crop-(old)crop introgressions? The overwhelming number of possible combinations between fully interfertile landraces and elite varieties of bread wheat, and the known lack of adaption to diverse agroecological conditions of most genotypes makes “random crosses and hoping for the best” a pre-breeding strategy with poor returns^1^. As an alternative approach less reliant on fortuity, we implemented a pre-breeding strategy based on wheat hybrids^3^ that predicts the respecting breeding value of PGR from a small training set of crosses (**Fig. 4**). To overcome the strong yield penalties suffered by non-adapted landraces (**Fig. 4a-b**), we estimated breeding values of first-generation ‘Elite PGR’ hybrids. Seven-hundred seven IPK accessions (**Supplementary Table 29**), pre-selected for high pollen shedding and synchronized flowering time, but retaining molecular diversity (**Fig. 4d** and **Supplementary Table 30**), served as pollinators for 36 different elite cultivars in an incomplete factorial mating design to obtain 1,427 Elite PGR hybrids (**Supplementary Fig. 9**). Multi-environment field trials established the high heritability of grain yield in such hybrids (**Supplementary Tables 31** and **32**). As expected, hybrids outperformed their PGR parents by a considerable margin (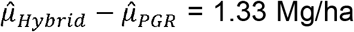, −log_10_(p-value) = 50.2, **Fig. 4b**). *Per se* performance of PGR was only weakly correlated with breeding value estimates (*r* = 0.22, p-value < 0.01), vindicating the expenses of hybrid seed production. We proceeded by developing ‘Elite PGR’ populations with PGR parents picked from the highest yielding (1^st^ decile) accessions according to their estimated breeding value in a hybrid background. After two-stage selection for appropriate height and good health, a substantial proportion (40%) of PGR-derived F_3:4_ families already outyielded the oldest (released in 2007) cultivar check in our yield trials (**Fig. 4c**, **Supplementary Tables 33** and **34**). Among the 10% superior families, TRI 4589 - an Uruguayan accession from the late 1950s - was the second most frequent PGR in the tested pedigrees, which suggests that even before breeding selection our strategy can highlight the contribution to future yield increases by abandoned, non-adapted PGR. The yield of the best family exceeded the average yield of check cultivars by 0.3 Mg/ha (4.4% improvement). The current annual genetic gain in wheat breeding in Germany is 0.7-1.2%^40^, indicating that our strategy achieved an impressive contribution to wheat improvement in less than one breeding cycle.

**Fig. 4.**
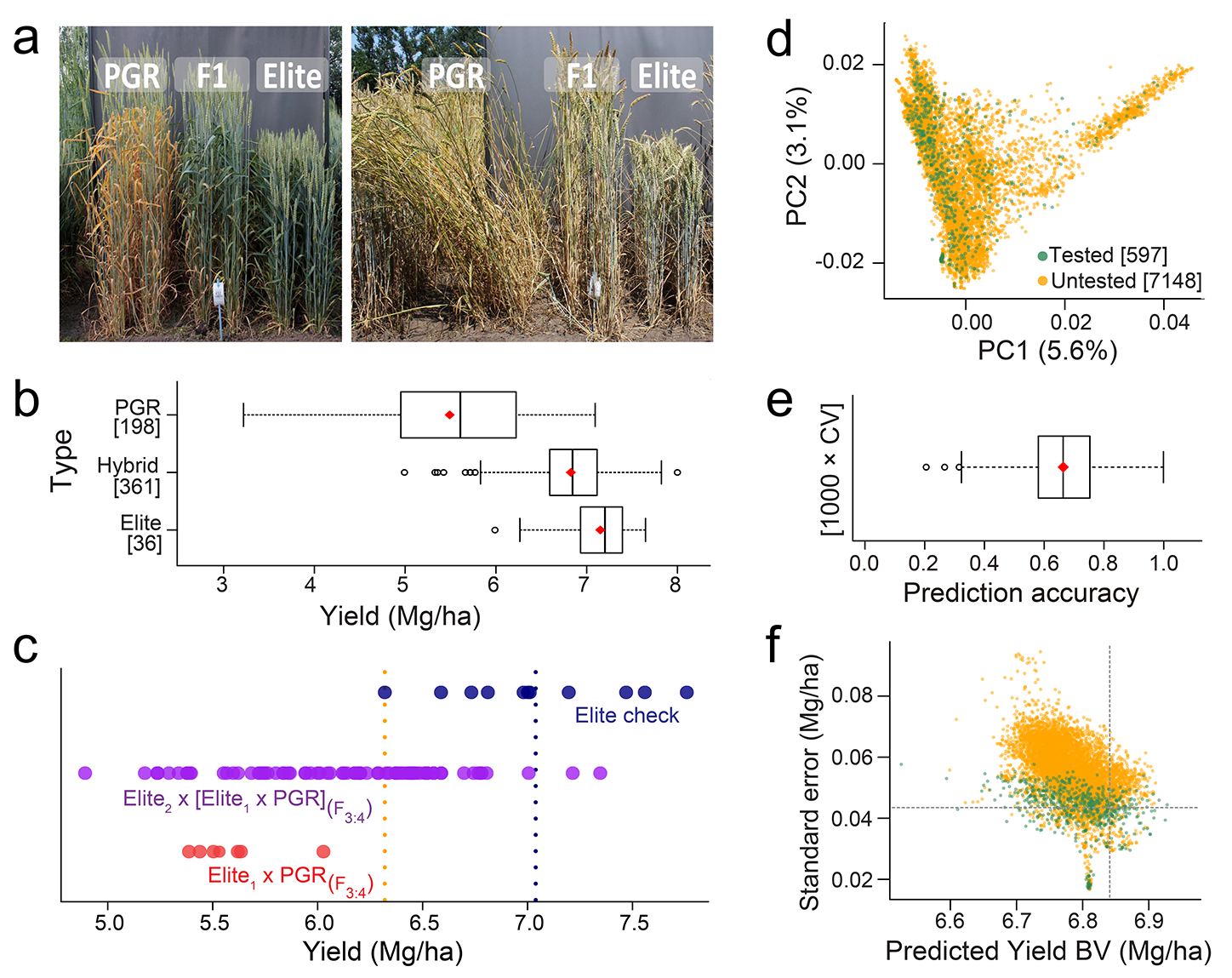
Uncovering the yield breeding value (BV) of plant genetic resources (PGR) for pre-breeding through ‘Elite×PGR’ hybrids and genomic prediction. **a**, Disease susceptibility (left) and lodging (right) mask the yield BV of wheat PGR, but these limitations are overcome in Elite × PGR (F_1_) hybrids, where PGR contribution to yield can be revealed. **b**, Parenting effect of elite cultivars on the yield (Mg/ha) of PGR observed in Elite×PGR hybrids. **c**, Yields of modern check cultivars as well as 85 F_3:4_ lines derived from two-(Elite_1_×PGR) and three-way (Elite_2_×[Elite_1_×PGR]) crosses with PGR parents selected for their high yield BV estimates. The performance of the oldest check and the average of checks are indicated with orange and blue vertical dashed lines, respectively. **d**, Molecular diversity stored at the IPK genebank as potrayed by the first two principal components (PC) from genotyping-by-sequencing variants and covered by yield BV estimates (green). These estimates were the training set for genomic prediction (GBLUP) of untested PGR (orange) using 29,844 SNP markers. **e**, Prediction accuracy of GBLUP in 1000 cross-validation (CV) runs. **f**, Yield BV genomic predictions and their standard errors (SE) for the IPK winter wheat collection. The vertical dashed line indicates the 10% superior yielding predictions while the horizontal one is the average SE of the training set. Red diamonds in (**b**) and (**e**) indicate the averages, while the numbers in brackets [ ] in (**b**) and (**d**) indicate the number of datapoints for each category.

A remaining uncertainty of our approach is whether our training set, whose size is constrained by the cost of hybrid seed production, is large enough to fuel robust genomic predictions. If so, the pool of possible donors – all accessions with yield predictions from cheap markers – would be greatly enlarged. We use genomic prediction to infer the breeding values for the entire IPK collection based on 29,844 SNP markers and using 597 PGR estimates from the hybrid context as a training set (**Fig. 4d-f**). The cross-validated prediction accuracy was high (0.66±0.12, **Fig. 4e**). Within the 10% superior fraction of the IPK genebank (> 6.84 Mg/ha), most predicted material (85.8%) has not been phenotyped and only 12.8% of these fully predicted high-yielders had standard errors (SE) as low as the average SE of the training set (**Fig. 4f**, **Supplementary Table 35**). High-yielding, highly reliable donors were predominantly from Europe (78.8% of 151 donors), followed by the Americas (4%) and Iran (1.3%) (**Supplementary Table 36**). Most these donors (66.2%) were acquired by the genebank before 1980, supporting our hypothesis that GiPS can contribute to rescuing exotic and past crop diversity.

Next, we predicted breeding values across the IPK and INRAE genebanks using IPK-PGR as training set (**Supplementary Fig. 10**, **Supplementary Tables 37** and **38**). The small number of shared markers affected the cross-validated prediction accuracy only mildly (0.60±0.13, **Supplementary Fig. 10b**). Out of 111 predicted high-yielding accessions with good statistical support, only six (5.4%) belonged to the INRAE collection (**Supplementary Fig. 10c** and **Supplementary Table 38**). All these INRAE donors originated in Europe, which is also the most common (71.1%) continent of origin for winter wheats stored at the INRAE genebank^20^. Importantly, none of these identified INRAE donors was part of INRAE’s unique diversity (**Supplementary Table 8**), which highlights that reliable predictions are only obtained within the diversity sampling space covered by the training set^16^. Among INRAE donors, five were registered as cultivars before 1970 and one of them even during the 1910s. In summary, GiPS can expedite the pre-breeding for a highly complex, and economically most relevant trait, grain yield, without discriminating against non-adapted germplasm.

## Discussion

The challenges associated with the use of genebank material in breeding and agriculture has been recognized for a long time^1^. We implemented a generally applicable strategy to close the gap between genebank management and pre-breeding. We have established the feasibility of selecting high-yielding donors from a small training set with yield records and dense marker data for a much larger universe of genebank accessions. The linchpin of GiPS are statistically robust yield estimates from early-generation Elite x PGR hybrids, which underpinned the statistically robust and meaningful inference of breeding values from a training set of hundreds of genotypes in thousands of accessions, even across genebanks. It goes without saying that all steps, selection of coresets, genomic prediction, and cross-referencing between genebanks, are dependent on genebank genomics^6^ and ultimately cheap genome sequencing. Unable to select donors only based on their marker profiles, two breeders’ professional life times would have been spent, if not wasted, in producing and trialing 7,000 ‘PGR x Elite’ hybrids.

Our analysis of current elite cultivars has shown that breeders did a good job by pyramiding large-effect resistance loci. In the absence of yield penalties linked to alien chromatin, breeders should go doing so^41^. Recent gene isolation efforts have focused on wheat wild relatives, notably its diploid wild D-genome progenitor, *Aegilops tauschii*^42^. Our genome-wide association has shown that also bread wheat proper harbors resistance genes, which after their isolation and functional characterization may become part of transgene cassettes. In contrast to *Ae. tauschii*, wheat landraces have a base-level adaptation to agricultural habitats that will likely facilitate the recombination of their haplotypes in breeding programs under jurisdictions unfavorable to genetic transformation.

Plant health is key to crop performance. Luckily, genetic architecture of plant resistance is relatively less-complex, with the main challenges being the pathogens overcoming major-effect genes deployed in isolation^28,43^. GiPS is a perfect complement to resistance gene stacking. Our results for grain yield, arguably the genetically most complex of traits, illustrate that the informed selection of donors is possible even if a complex genetic architecture and pervasive genotype-by-environment interaction prevent singling out causal genetic factors.

## Methods

### Plant material and defined seed for IPK-PGR

The winter wheat plants in the current work trace back to 9,145 PGR from the IPK-genebank and 337 diverse approved elite cultivars (cultivars panel) plus 131 diverse elite inbred lines (breeder’s panel). Their passport data regarding acquisition (PGR) and release (elite cultivars) year, geographic origin, as well as growth habit is presented in **Supplementary Table 1**. Origins were further grouped into 14 different macrogeographic regions according to Balfourier et al.^20^. Passport data for PGR registered as TRI number were accessed via the Genebank Information System^44^ (GBIS) in extended MCPD-format. Passports of PGR from the B number register and of elites were compiled from various databases and publication sources. A total of 55 different geographic origins were reported for PGR, with Germany (14.2%), followed by Italy (8.1%), countries of the former Soviet Union (6.5%) and the USA (6.3%) being the most common origins. Europe is the most common macrogeographic region for PGR (60.6%), with the greatest proportion (38.3% of European PGR) coming from Western Europe. South Asia and North America are represented by 9.5% and 6.6% of PGR, respectively. Most PGR (97.8%) were acquired before the 2000s, with the oldest one tracing back to 1927 and the newest acquisitions to 2007. The cultivar panel was almost entirely composed of winter types (95.5%) plus some facultative or spring types. Breeders or commercial owners can be traced to ten different European countries, with most cultivars released by Germany (47.2%), followed by Great Britain (15.4%), France (14.2%), and Poland (9.8%). Cultivar release dates ranged from 1975 to 2018, with most cultivars (82.5%) released from 2000 onwards. ‘Monopol’, the oldest cultivar, was released in Germany during 1975 and is still grown today for its high milling and baking quality (https://ig-pflanzenzucht.de/sorte/monopol/). The panel’s most recent material corresponds to the German cultivar ‘Informer’ released in 2018. The German inbred lines belonging to the breeder’s panel were sampled from 4 breeding companies in the first half of 2010s^45,46^.

Within the single-row multiplication plots, one representative ear was bagged for homogeneous plots, while up to two ears were selected for clearly non-homogenous PGR. Seeds from isolated ears were harvested separately from the rest of the plot and further propagated using an ear-to-row method. Plant material from isolated ears is referred to as SSD-PGR to differentiate them from the accessions (PGR). A digital object identifier (DOI) was assigned to each SSD-PGR whose PGR donor was registered in GBIS (**Supplementary Table 39**). Seeds for the elite cultivars were sourced from local markets, while the breeder panel was supplied by four German-based breeding companies^45,46^.

### Library Preparation and sequencing

For DNA extraction, ten seeds of one genotype were grown in the greenhouse and a single, approximately 10 cm leaf was harvested from a 10-days-old seedling. DNA extraction was performed using a silica-membrane technology (NucleoSpin® 96 Plant II) as described by the manufacturer (Machery-Nagel). A total of 7,745 SSD-PGR and 325 genotypes from the cultivars panel were characterized using GBS following the protocol for digestion with two restriction enzymes^47,48^. For this, DNA samples were simultaneously digested with *Pst*I and *Msp*I (New England Biolabs) and ligated with adapters containing sample-specific barcode sequences. Later, the processed barcoded DNA samples were pooled into groups of 540 genotypes in equimolar amount to form a GBS library. Single-end sequencing (1 × 107 cycles) was performed on Illumina Hiseq-2500 or NovaSeq 6000 system using custom sequencing primers according to manufacturer’s instructions (Illumina). WGS was carried out on 263 diverse SSD-PGR from the T3Cs plus 191 diverse genotypes from the cultivar panel (**Supplementary Note**), and for the 131 elite lines of the breeder’s panel. The SSD-PGR fraction was extended with further WGS data for 183 PGR from the GenDiv-Project. WGS libraries were prepared using the Nextera DNA Flex Library Prep according to the manufacturer’s (Illumina) instructions. Libraries were pooled in an equimolar manner. The multiplexed pool was quantified by qPCR and sequenced (paired-end, 2 x 151 cycles and 10 bp for the index reads) using a NovaSeq 6000 device (Illumina) at 3-fold coverage.

### Read processing and Variant calling

For GBS, the adapter sequences and low-quality bases from raw reads were trimmed using cutadapt (v1.16)^49^ with a minimum read length of 30 bp. Adapter and quality trimming was further confirmed by using FastQC^50^. The high-quality reads were aligned against the hexaploid wheat reference genome assembly cv. Chinese Spring (RefSeq v1.0)^22^ using BWA-MEM (v0.7.17)^51^ with default parameters. The output was converted to binary alignment map (BAM) format using SAMtools (v1.9)^52^ and then sorting was performed using NovoSort (v3.06.05). Variant calling was performed using the mpileup and call functions from SAMtools (v1.9) and BCFtools (v1.8)^52^. The software was run with -DV parameter for SAMtools mpileup and minimum read quality (q) cutoff of 20. The bi-allelic SNPs were further filtered with minimum QUAL ≥ 40, minimum read depth for homozygous call ≥ 2, and minimum read depth for heterozygous calls ≥ 4 using a custom awk script. The vcf files were imported into the R statistical environment^53^ (v3.4.3) and converted to GDS format for further processing using seqArray^54^.

For WGS, variant calling was performed as mentioned above except that minimap2^55^ was used for read alignment. The minimap2 was run with genome index size of 50Gb and keeping other parameters to default. The bi-allelic SNPs with minimum QUAL ≥ 40, minimum read depth for homozygous call ≥ 1, and minimum read depth for heterozygous calls ≥ 2 were imported into the R statistical environment and filtered as mentioned above. All post-filtering criteria regarding missing values, homozygous and heterozygous calls, of GBS and WGS-SNP data used in the different analyses are summarized in **Supplementary Table 2**.

### Genetic diversity within the IPK collection

PCA was performed using the snpgdsPCA() function of SNPRelate^56^, which implements a FastPCA algorithm^57^. The population structure analysis was carried out using model-based clustering approach implemented in ADMIXTURE^58^. The software was run for different K values from 2 to 15 with 10-fold cross-validations (CV) and 500 bootstrap replicates. Fst between populations were calculated using vcftools^59^. Furthermore, the identity-by-state (IBS) analysis was performed using the snpgdsIBSNum function in the R statistical environment. The proportion of pairwise difference (PPD) between two samples was calculated using the formula: IBS0/(IBS0+IBS2). All pairs with PPD value ≤ 0.01 were selected and clustering based on PPD values was performed to identify nearly identical samples (duplicates) using the R package igraph^60^ for graph operations.

### Comparison of genetic diversity between IPK and INRAE genebanks

A sample of 2,608 PGR from the INRAE winter wheat collection was previously genotyped^20^ using a high-density Affymetrix Axiom SNP array^61^. To compare diversity between IPK and INRAE genebanks, SNPs from the array were mapped to RefSeq v1.0 and their positions extracted. With a threshold for missing values (< 50%) in GBS-SNPs from the IPK collection, a total of 895 SNPs were retained in the merged data set. PPD values were not only used to identify duplicates between genebanks but additionally, to detect diversity unique to each genebank. An accession was considered as unique to its genebank when its minimum genetic distance to all accessions of the other genebank was greater than the 95% quantile of distances within the IPK collection.

### Introgression identification and tracing the history

A *k*-mer-, i.e. unique sequence of n bases, based approach was used to describe the genotypes for previously known introgressions^6^. The sequences of introgressed regions were extracted from the chromosome scale assembly of respective genotypes^9^. The KMC tool (v3.0)^62^ was used to identify 71 bp unique *k-*mers from each introgressed sequence separately. The 71 bp *k*-mers were also identified from RefSeq v1.0. The *k*-mers from each introgressed region were subtracted from RefSeq v1.0 *k-*mers using the KMC tool to identify unique and specific *k*-mers for each introgressed region. The *k*-mers were then compared with quality-checked short reads for each genotype, and reads carrying *k*-mers were counted. The proportion of reads for each introgressed region was plotted against the total number of reads to identify genotypes with introgressions.

To identify new introgressions, the BAM files generated during the variant calling process were used. The trimmed reads from Chinese Spring were also mapped against RefSeq v1.0 and a BAM file was generated. From the BAM file for each genotype, the number of reads in the 500Kb window were calculated. Reads from each window were first normalized to sequencing coverage and then to the number of reads from the same window for Chinese Spring. The logarithm of normalized count was plotted to create a genome-wide coverage plot.

### Nucleotide diversity and selective sweep study

To identify regions under selection, the WGS nucleotide diversity was examined within populations defined by their release/acquisition periods: 255 genotypes from the Green Revolution (PreGreen), 212 varieties released between 1971 and 2000 (OldCV), and 293 recent genotypes bred after 2000 (NewCV). Nucleotide diversity analysis was carried out for all possible pairwise population combinations. Normalized XP-CLR^24^ scores for each combination were ordered in descending order and the top 0.1% intervals were interpreted as regions under selection (selective sweep). Regions that were 10Kb adjacent were merged. Nucleotide diversity analyses were conducted using pixy^63^ with a 10Kb window and a step size of 100 bp, while leaving all other parameters at default.

### Phenotyping for yellow rust resistance

YR screenings were based on naturally occurring infections for 7,684 PGR along with 80 elite checks in 12 unbalanced, replicated field experiments considering 1,428-1,697 entries each (for further details see **Supplementary Table 10**). On average, each PGR was tested in 2.4 experiments, with almost all (99.4%) tested in at least 2 different experiments. Elite checks were used to increase the connectivity between experiments, with each check being present in 5.1 experiments on average, while four checks were present in all experiments. Field experiments were conducted between harvest years 2015-2020 considering locations Gatersleben and Schackstedt. An alpha lattice design with two complete replications divided into incomplete-blocks was used to account for uncontrolled spatial variation in each field experiment. Following the official protocols of the German Federal Plant Variety Office^64^, an ordinal scale from 1 to 9 was used to score infections, where one stands for minimal symptoms and nine indicates extensive disease symptoms.

With the goal of finding new sources of disease resistance in PGR that are lacking in elite breeding, T3C for three leaf diseases were assembled (**Supplementary Note**). Briefly, we used preliminary phenotypic data and performed an unbalanced two-tailed selection to enrich the fraction of resistant genotypes while selecting susceptible genotypes such that the association between population structure and the trait under consideration is minimized. The complete set consisted of 200 genotypes from the cultivars panel and 600 SSD-PGR from the three T3Cs (**Supplementary Table 15**).

YR screenings were then performed for the diverse elite cultivars and the T3Cs in an additional set of 7 balanced experiments (further details in **Supplementary Table 16**). Briefly, experiments were conducted during harvest years 2019 and 2020 in locations Gatersleben, Quedlinburg, Wetze and Rosenthal. The two experiments in Gatersleben relied on natural infections while the other five were artificially inoculated. The percentage of YR infection from the experiment in Quedlinburg was transformed into a 1-9 scale as specified in **Supplementary Table 40**. For the other six experiments, infection was scored using the 1-9 ordinal scale as previously outlined.

### Estimating breeding value for grain yield in a hybrid background

Individual PGR often lack some major loci for adaptation to modern agricultural practices. This masks their breeding value when evaluated in yield trials. As a rapid adaptation strategy, it has been proposed to estimate the breeding values of PGR in hybrid backgrounds when crossed with elite cultivars^3^. This strategy was implemented in five consecutive years for a total of 760 PGR (year 1 PGR, year 2-5 SSD-PGR) by crossing each of them as male parents with up to 14 elite cultivars, with an average of 2.1 per PGR. To ensure a sufficient quantity of hybrid seed, PGR were selected based on their pollination capability. The details of how hybrid seed production was carried out and also the phenotyping of series one to three have been described in detail previously^65,66^. Across-series, a total of 22 unbalanced field experiments were conducted to evaluate grain yield of 1,925 hybrids, along with a set of 518 parental lines and 40 checks to improve connectivity between series. Unbalanced experiments span together harvest years 2016-2020 and seven different German locations. Details of the experimental designs for each experiment series can be found in **Supplementary Table 31**.

### Using breeding value estimates of PGR for an informed parent selection

Based on the field trials conducted in the first year, we embarked on a pre-breeding program that used the estimated breeding values for grain yield as a tool for PGR parent selection. We applied stringent selection with a superior fraction of 10% and developed segregating populations using two-(Elite_1_×PGR) and three-way crosses (Elite_2_× [Elite_1_×PGR]). The segregating progeny were genetically fixed by two generations of selfing. Simultaneously, we performed two-stage selection based on visual assessment of single plants, followed by rows focusing on the traits plant height and leaf health. In 2020, 85 advanced F_3:4_ families were evaluated for yield under conventional local agricultural practices at two locations and considering eleven winter wheat cultivars approved for commercial use in Germany during the last one and a half decades as checks (**Supplementary Table 33**). Among checks, the French cultivar ‘Arezzo’ (released in 2007) was the oldest one while the German variety ‘Informer’ (released in 2018) was the most recent cultivar (**Supplementary Table 1**). In these experiments as well as for the 22 unbalanced field experiments with ‘Elite PGR’ hybrids, plots were mechanically harvested while grain yield was expressed in Mg/ha on a 140 g H2O per harvested kg moisture basis.

### Phenotypic data analyses of yellow rust screenings

Outlier detection, BLUEs and variance components for single replicated experiments were obtained using equation (4) of the **Supplementary Note**. Regarding unbalanced experiments (**Supplementary Table 10**), the YR infection per plot in Gatersleben during 2019 was the maximum value among early and late scorings for these analyses, while the unreplicated Schackstedt experiment from 2019 was only considered for analyses across-experiments. Variance components and BLUEs across unbalanced experiments were obtained in an integrated outlier-corrected dataset using equation (5) of the **Supplementary Note**. For the elite cultivars plus T3Cs (**Supplementary Table 16**), each YR scoring date was analyzed separately and the unreplicated data from Quedlinburg were corrected by subtracting the incomplete-block means. Later, the maximum outlier-and-design-effects-corrected value among early and late scorings was selected entry-wise in each experiment and integrated across-experiments. Across-experiments BLUEs and variance components for the elite cultivars plus T3Cs were obtained through the following mixed model:

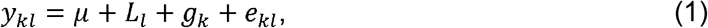

where *y_kl_* is the YR infection value for the *k*-th genotype in the *l*-th experiment, μ corresponds to the general mean, *L_l_* denotes the effect of the *l*-th experiment, *g_k_* indicates the effect of the *k*-th genotype, while *e_kl_* is the error term of the model
confounded with the genotype experiment interaction. Assumptions for BLUEs and variance components computation are outlined in the **Supplementary Note**. A pooled error variance 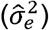 was obtained by averaging error variance estimates of single experiments analyses. Later, the variance estimate of experiment × genotype interactions 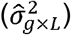 was obtained as 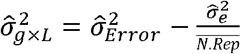, where 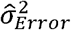 is the error variance estimate from equation (1) and 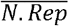 is the average number of replicates across experiments. Estimated variance components were used to compute heritatabilities for single and across-experiments according to equations (6) and (7), respectively (**Supplementary Note**).

### Estimating breeding value for grain yield of PGR

Previously published procedures for data curation and estimation of genotypic effects corrected for experimental design effects using linear mixed models were implemented^65,66^. The basic model included the effects of genotypes and incomplete-blocks in addition to other experiment-specific (trials and/or replications) design effects (**Supplementary Table 31**). All data were checked series-wise for outliers using Method 4 “Bonferroni-Holm with rescaled residuals standardized to mean absolute deviation” described by Bernal-Vasquez et al.^67^. Outliers were removed and the linear mixed model described above was refitted to obtain estimates of genotypes in each experiment, which were adjusted for the effects of experimental designs and served as input to subsequent analyses. In a next step, 161 hybrids with low seed purity were discarded from the integrated analyses. Adjusted means for genotypes from the 22 experiments were used in a linear mixed model including effects of groups - i.e., genotypes belonging to either checks, lines, or hybrids -, experiments, lines, breeding values of PGR and elite lines, i.e. the general combining ability effects, their interaction effects with experiments, specific combining abilities effects and errors. Assuming all model effects excepting group means as random, variance components and PGR breeding values were estimated. Variance estimates were subsequently used to compute the broad-sense heritability of the hybrid yield performance as specified in Zhao et al.^66^, while a narrow-sense heritability of the PGR contribution to yield in the hybrid context was obtained as follows:

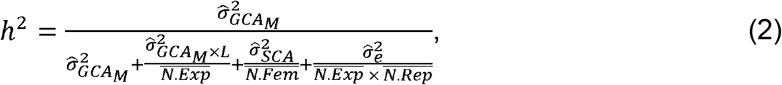

where 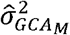 is the variance estimate of the general combining abilities of male parents, 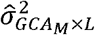 denotes the variance of interactions between male combining abilities and experiments, 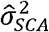 indicates the variance of specific combining abilities between parents, 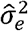 corresponds to the error variance, while 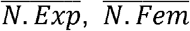 and 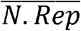 are the average number of experiments, crossing females and replications.

### Genome-wide association study (GWAS)

Unimputed GBS-SNPs were used to compute a kinship matrix as 2 × (**J**-**RD**), where **J** denotes an n×n all-ones matrix and **RD** is a Rogers’ distance^68^ matrix calculated as stated in the **Supplementary Note**. Unimputed WGS data were used to conduct GWAS for YR. GWAS was performed using a standard linear mixed model^69^ that corrects for population stratification by fitting a polygenic effect for the genotypes, with covariance modeled using the previously estimated kinship matrix. For genome-wide multiple test correction, the nominal significance level 0.05 was divided by the number of independent markers (*M_eff_*)^70^.

In addition, we implemented a GWAS approach which potentially captures structural variation denoted as *k*-mer based GWAS^71^. Briefly, KMC tool (3.1.1)^62^ was used to identify 31 bp *k*-mer from quality trimmed sequencing reads from each genotype. Only *k*-mer supported by at least 2 reads were considered. *k*-mer from all the genotypes were combined to generate a non-redundant *k*-mer set. The non-redundant *k*-mer sequences were searched in each genotype and a *k*-mer presence-absence matrix was generated which served as a genotype matrix in GWAS. To reduce computational burden of GWAS for billions of *k*-mers, we selected the first top 10 million *k*-mers based on the *T*^2^ statistic as implemented in the kmers_gwas.py script^71^. Afterwards, GWAS was conducted for this reduced set of k-mers as stated before for SNP-based GWAS but using a Bonferroni significant threshold for multiple-test correction. The *k*-mers were mapped against reference genome sequences using BWA-MEM and only uniquely mapped *k*-mers were extracted. The linear mixed models of phenotypic analyses as well as GWAS were fitted using the average information matrix algorithm for restricted maximum likelihood (REML) computation as implemented in ASReml-R (3.0)^72^.

### Genome-wide predictions

Genomic best-linear unbiased prediction (GBLUP) was performed with an additive genomic relationship matrix according to the first VanRaden method^73^. Predictions were obtained for the 7,745 SSD-PGR as well as for the INRAE-PGR by using a training set of IPK-PGR having both phenotypic and genotypic data. Missing values still present after filtering were imputed using the locus mean. In addition, the standard error (SE) of prediction for each genotype *i* was calculated as 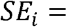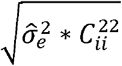, where 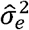 is the genomic-estimated error variance and 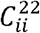 is the *i* diagonal element of the lower-right quadrant from the generalized-inverse of the left-hand-side coefficient matrix according to Henderson^74^. The expected prediction accuracy was estimated through five-fold cross-validation within the training set and was defined as the Pearson correlation coefficient between predicted and observed values divided afterwards by 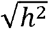. Mixed model equations for genomic prediction were computed using REML as implemented in the R-package rrBLUP^75^. All computational methods related to phenotypic analyses, GWAS, and genomic prediction were implemented within R statistical environment^53^ (v3.6.1).

## Acknowledgements

This research work was mainly funded by the German Federal Ministry of Education and Research under the frame of the Project GeneBank2.0 (grant no. FKZ031B0184B and FKZ031B0184A). Additional financial support was provded by the German Federal Ministry of Food and Agriculture under the frame of the GenDiv-Project (grant no. 2814603813). Authors are very thankful to Dr. Andreas Börner for providing seeds of the B catalog. Authors would like to also thank Christoph Martin, Jelena Perovic, Johannes Schneider, Sonja Gentz, Andrea Kunze, Martina Kühne, Lena Gaczensky and Martin Koch for their valuable technical support in field activities, as well as Susanne König, Jacqueline Pohl, Ines Walde and Manuela Knauft for their technical assistance in producing GBS and WGS data. Authors additionally thank Jens Bauernfeind, Thomas Münch and Heiko Miehe for administration of the IT infrastructure as well as Anne Fiebig, Danuta Schüler and Daniel Arend for their support in the data publication.

## Author contributions

J.C.R., M.M., N.S. and V.K. developed the concept. M.O. and S.W. provided passport information of the TRI catalog of genebank accessions as well as DOIs for their derived progenies. S.K. provided DNA for GBS and A.H. and N.S. produced sequencing raw reads. A.H. and N.S. obtained high quality DNA samples and generated WGS raw reads. A.W.S., N.Philipp, U.B., A.S., N.Pfeiffer., P.H.G.B. and J.S. conducted YR resistance screenings. N.Philipp, P.H.G.B. and C.F.H.L. produced seed and conducted yield trials for hybrids. N.Philipp, M.R. and J.C.R. produced, selected and yield-tested PGR-derived families. J.F. confirmed wheat ploidy level through fluorometry. A.W.S., Y.Z., N.Philipp and M.R. analyzed and curated phenotypic data. S.M.K. processed sequencing reads, integrated INRAE and IPK genomic data, generated SNP and *k*-mer matrices and performed diversity, selective sweep as well as introgression analyses. A.W.S. integrated genomic and phenotypic data, selected T3Cs and performed genomic prediction. F.L. performed GWAS for YR and selected donors with the support of J.C.R., A.W.S. and M.M.. Y.J., Y.Z. and M.M. provided statistical support. M.L. and U.S. facilitated the data management, the sequence and variation data submission to public repositories. A.W.S., S.M.K., F.L., M.M. and J.C.R. wrote the manuscript with the input of all other co-authors.

