## Supplementary note and figures for "GiPS: Genomics-informed parent selection uncovers the breeding value of wheat genetic resources"

The goal was to assemble a core collection that would enable the discovery of new sources of resistance against *Puccinia striiformis* f. sp. *tritici* a.k.a. yellow rust (YR) in wheat plant genetic resources (PGR) from the German Federal *ex situ* Genebank for Agricultural and Horticultural Crop Species (IPK-genebank) through association mapping. The following describes the strategy we used to establish a trait-customized core collection (T3C) based on preliminary data for YR resistance available at the beginning of the study.

***Generation of preliminary yellow rust resistance screening data to select the T3C***

Seven series of preliminary experiments, i.e., A to G, were conducted to screen wheat PGR from the IPK-genebank for YR resistance (**Supplementary Table 41**). The experiments were conducted at a total of three locations between 2014 and 2018. While experimental series A was divided into five single experiments, each experimental series B to G corresponds to a single experiment itself. Experimental series A is subsample of the 12 unbalanced experiments performed to screen a total 7,684 PGR between 2014 and 2019 crop seasons (**Supplementary Table 10**). The number of entries tested within a single experimental series ranged from 144 (series G) to 3,073 (series A). With the exception of experimental series B, in which resistance screenings were conducted with seedlings grown in pots under greenhouse conditions, all other experiments were performed in the field, where YR severity was measured in plots (0.2-0.4 m^2^) at the adult plant stage. In addition, experimental series B was the only one where entries were not replicated and seeds of the tested PGR were traced back to the single ear(s) isolated in multiplication plots (SSD-PGR). For the rest of the series, the seeds of each PGR corresponded to one sample per accession. While the screenings of experimental series B to F were performed with a completely randomized design, the experiments from series A and G followed an alpha lattice design with incomplete blocks. YR infections of experimental series A, C, D, and G were based on natural infections, whereas those in series B, E, and F were based on artificial inoculations with a YR isolate race mixture collected from previous wheat seasons. Experimental series C and D were originally artificially inoculated to screen for resistance against *Puccinia triticina* f. sp. *tritici* (leaf rust, LR) and *Fusarium culmorum*, respectively, but were also considered due to the presence of naturally occurring YR. For the greenhouse experiment (series B), symptoms of leaf infection were scored at a single time point, ten days before 10-days-old moist seedlings were sprayed with a liquid suspension of white clay powder and YR inoculum. In the field experiments artificially inoculated with YR (Series E and F), infected adult plants of “spreader plots” sown with known susceptible genotypes served as the initial inoculum source for the tested material. Here, leaf infection symptoms for each plot were scored and recorded once a week for four consecutive weeks starting ~1 month after inoculation. In experimental series A, C, and D, symptoms were scored only once for each plot after sufficient disease pressure of naturally occurring YR was observed for entire experiments. In experimental series G, YR leaf symptoms were evaluated once a week for each plot for three consecutive weeks after a general natural YR outbreak was observed for the entire experiment. The rating scale for experimental series B was qualitative^76^, while an ordinal rating scale from 1 to 9 according to the Protocols of the German Federal Plant Variety Office^64^ or a percentage scale was used for the remaining experimental series. To make the results of different experimental series more comparable, all measurements were transformed into a percentage scale using the key presented in **Supplementary Table 40**. After transformation, either the maximum score among multiple measurements or the score of a single measurement time of a pot/plot was used for further analyses. Exclusively for experimental series G, the maximum values of each original and transformed percentage scale were averaged for each plot and used for further analyses.

***Phenotypic data analyses of preliminary yellow rust screening experiments***

Different mixed models were applied according to the design of replicated experiments. For experimental series C to F, the following model was fitted experiment-wise according to the complete randomized design:

$y_{jk(l)}=\mu_{(l)}+g_{k(l)}+e_{jk(l)}$,	(3)

where $y_{ijk(l)}$indicates the YR infection value of the $j$-th replication for the $k$-th entry tested in the $l$-th experiment, $\mu_{(l)}$ denotes the common mean of the $l$-th experiment, $g_{k(l)}$ corresponds to the effect of the $k$-th entry, and $e_{ijk(l)}$ is the error term of the model.

For experimental series G as well as for the five single experiments within experimental series A, equation (3) was extended to accommodate effects of the alpha lattice design and fitted experiment-wise as follows:

$y_{ijk(l)}=\mu_{(l)}+r_{i(l)}+b_{j(il)}+g_{k(l)}+e_{ijk(l)}$,	(4)

where $y_{ijk(l)}$is the YR infection value for the $k$-th entry tested in the $j$-th incomplete-block within the $i$-th replication of the $l$-th experiment, $\mu_{(l)}$ and $g_{k(l)}$ are as previously defined in equation (3), $r_{i(l)}$ denotes the $i$-th replication effect, $b_{j(il)}$ is the $j$-th incomple-block effect nested within the $i$-th replication, and $e_{ijk(l)}$ accounts for the error term of the model. Due to the vast amount of data points for experimental series A (YR scores for 15,215 plots), an outlier test was performed experiment-wise following Anscombe and Tukey^77^ by assuming $\mu_{(l)}$ and $g_{k(l)}$ as fixed and the remaining factors in equation (4) as random. After removing datapoints flagged as outliers, the following model was fitted to the outlier-corrected experimental series A:

$y_{ijkl}=\mu+L_{l}+r_{i(l)}+b_{j(il)}+g_{k}+L_{l}\times g_{k}+e_{ijk(l)}$,	(5)

where $\mu$ is the mean across experiments, $L_{l}$ is the main effect of the $l$-th experiment, $g_{k}$ is the mean entry effect across experiments, $L_{l}\times g_{k}$ is the interaction between the $l$-th experiment and the $k$-th entry, while the remaining terms of the model retain all previous definitions of equation (4).

For the computation of the best linear unbiased estimations (BLUEs) of entries, the common mean and entry effects in equations (3) to (5) were assumed as fixed while the rest of the terms were considered random. BLUEs from each experimental series as well as YR infection scorings of experimental series B were used to calculate the correlation between overlapping entries. By assuming all factors excepting the common mean in equations (3) and (4) as random, variance estimates of entries ($\hat{\sigma}_{g}^{2}$) and errors ($\hat{\sigma}_{e}^{2}$) were computed. In case of experimental series A, these estimates were computed experiment-wise using outlier-corrected datasets. In addition to $\hat{\sigma}_{g}^{2}$ and $\hat{\sigma}_{e}^{2}$, the variance estimate of the interaction between entries and experiments ($\hat{\sigma}_{g\times L}^{2}$) was computed for the experimental series A by assuming all factors excepting the common mean in equation (5) as random. Heritabilities were then derived from the estimated variance components as follows:

$h^{2}=\frac{\hat{\sigma}_{g}^{2}}{\hat{\sigma}_{g}^{2}+\frac{\hat{\sigma}_{e}^{2}}{\bar{N.Rep}}}$,	(6)

where $\bar{N.Rep}$ is the average number of replications used to test entries. For experimental series A, $h^{2}$ was computed as:

$h^{2}=\frac{\hat{\sigma}_{g}^{2}}{\hat{\sigma}_{g}^{2}+\frac{\hat{\sigma}_{g\times L}^{2}}{\bar{N.Exp}}+\frac{\hat{\sigma}_{e}^{2}}{\bar{N.Exp} \times\bar{N.Rep}}}$ ,	(7)

where $\bar{N.Exp}$ is the average number of experiments to test entries. All mixed model equations were fitted using ASReml-R (3.0)^72^ within R environment^53^ (v3.5.1).

***Description of processed preliminary yellow rust resistance screening data***

Briefly, phenotypic analyses provided quality measurements for YR resistance scorings in replicated experiments and allowed to reduce their complexity by correcting for design effects (**Supplementary Table 42**). Within experimental series A, only few (3.6 % at the most) of the plots within a single experiment were flagged as outliers and excluded from further analyses. With the exception of experimental series D, YR resistance scorings were moderate to highly heritable ($h^{2}$ ≥ 0.63). Low heritability in experimental series D were presumably caused by the confounding effects of the artificial infection with *Fusarium culmorum*. Moreover, the highest heritabilities ($h^{2}$ > 0.9) were estimated for experimental series E and F. The presence of negative BLUEs for YR infection scores is presumably due to an overcorrection of experimental design effects for entries whose infection levels were extremely close to zero. YR screenings of experimental series D and those of SST_2015_2 from experimental series A presented the lowest (1.5%) and highest (24.3%) average infections levels, respectively. The broadest range of YR severity amounted to 81.8% and was observed for the across-experiments BLUEs of experimental series A. In contrast, the narrowest infection range (16.4%) was found for entries tested in experimental series G. The highest number of overlapping entries (1,442) was observed between experimental series A and B (**Supplementary Table 43**). Due to the vast number of tested entries within each of these two experimental series, their overlaps with other experimental series were also maximized. On the contrary, experimental series G had in general the lowest number of overlapping entries with other experimental series and presented in the best case only 20 overlapping entries (experimental series A). Except for the low and moderate significant correlations of YR severity from experimental series F with those of series D and C, respectively, all other correlations were not significantly different from zero. Particularly, the lack of association between seedlings tested in the greenhouse (series B) and adult plants tested in field experiments (series A, C, D and F) reflects the different types of YR resistance mechanisms been targeted by these two types of experiments.

***Preliminary marker data available before core-selection***

Genotyping-by-sequencing (GBS) data were produced as outlined in the Methods section. A total of 283,720 SNPs with mac 34 and 95% missing data were selected. The missing data were separately imputed for elite and PGR samples pools using the FILLIN (Fast, Inbred Line Library ImputatioN) algorithm implemented in TASSEL^78^ (v5). The imputed matrices for 2,828 SSD-PGR as well as 322 elite cultivars were integrated into a single matrix that was filtered again to retain 33,341 SNPs having less than 5% missing values and MAF > 0.01. Afterwards, SNPs were coded as 0, 2, 1, for the first and second alleles at homozygous and heterozygous states, respectively.

***Integration of preliminary genomic and phenotypic data***

Initially, only those PGR tested in experimental series A for which both GBS profiles and a sufficient quantity of seeds from isolations were available were used for the integration of infection values across experimental series into a mean infection index ($MII$) in the way:

${MII}_{(k)}=\frac{1}{{PI}_{(k)}}\sum_{l=1}^{{PI}_{(k)}} {PS}_{(k)l},$	(8)

where ${PI}_{(k)}$ is the phenotyping intensity, i.e. the number of experimental series, for the $k$-th entry and ${PS}_{(k)l}$is the percentage score at the $l$-th instance for the $k$-th entry. For experimental series A, only the BLUEs across experiments were considered, while raw transformed seedling infections for experimental series B and single-experiment BLUEs for series C, D, E, F and G were considered for $MII$ computation.

Pairwise Rogers’ distances^68^ (RD) between entries were calculated based on the filtered SNP data. Considering the total number of entries as $n$ and the total number of loci as $p$, the matrix of SNPs can be represented as $\boldsymbol{M}\boldsymbol{=(}m_{is}\boldsymbol{)}$**,** where $1\leq i\leq n$ and $1\leq s\leq p$. Given $1\leq j\leq n$, the RD between any $i$-th and $j$-th entries is calculated in the way:

${RD}_{ij}=\frac{1}{2p}\sum_{s=1}^{p} |m_{is}-m_{js}|$.	(9)

RDs were compiled into an $n\times n$ RD matrix for downstream analyses.

***Using a T3C-strategy to allocate limited resources for intensive characterization and mine genebanks for novel functional variation underlying yellow rust resistance***

*1) Elite core.* To define the variation of PGR as novel, a reference point on the variation currently present in cultivars is needed. Therefore, a core-set of 200 elite cultivars having sufficient seeds to conduct intensive field testing was selected among 317 lines from the cultivars panel for which GBS data were available (**Supplementary Fig. 5**) by applying the maximized average entry-to-nearest-entry distance (EN) algorithm on the estimated RD. As implemented in the corehunter^79^ R-package (v3.2.1), this algorithm is expected to maximize neutral molecular diversity and minimize redundancy within selected cores.

*2) PGR cores.* Allocating the same number of slots for intensive characterization of PGR was more challenging than for elite cultivars. Due to uncertainties in seed multiplication, an unexpected 10% reduction in the number of SSD-PGR with available seeds was assumed. Thus, 220 instead of 200 available slots were considered during selection, and final numbers were reduced to 200 according to their seed availability after selection. A two tailed-selection was applied to select the PGR cores as follows: According to the calculated $MII$, SSD-PGR responses to YR are diverse, varying from 0% to 81.13% infection (**Supplementary Fig. 5b**). Contrasting phenotypic variation is essential for performing association mapping^80^; therefore, not only the 10% most resistant but also the 10% most susceptible fractions among the 2,486 SSD-PGR were selected first. Nevertheless, identification of repeatable or reliable sources of pathogen resistance requires intensive testing^81^. Because the $PI$ for PGR ranged from 1 to 5 experimental series, a cutoff of $PI$ > 2 was first applied to filter out reliable resistant PGR. Among the remaining SSD-PGR, the 249 most resistant ones were selected (**Supplementary Fig. 5b**). In contrast, no $PI$ filtering was performed for the 249 most susceptible SSD-PGR. Within the resistant tail, the $MII$ ranged from 0.38% to 13.6%, with median and mean values of 6.4% and 6.8%, respectively. For the susceptible tail, the $MII$ ranged from 27.5% to 81.1%, while the median and mean values here were 33.9% and 38.2%, respectively. SSD-PGR from resistant and susceptible tails were broadly distributed and covered to a large extent the neutral molecular diversity represented by the first two coordinates (PCo1 and PCo2) from a principal coordinate analysis of the RD matrix using the cmdscale() function (**Supplementary Fig. 5c**). Since the focus was to find new sources of YR resistance, ¾ of the formerly available resources were allocated to resistant SSD-PGR, while the remaining ¼ were for the susceptible ones. Therefore, the sizes of resistant and susceptible tails were reduced in the following to 165 and 55 SSD-PGR, respectively.

In addition to maximizing diversity and reducing redundancy, the 165 SSD-PGR from the resistant tail were selected based on their ability to maximize PGR diversity that was not covered by the elite cultivars (**Supplementary Fig. 5d**). To this end, the previously selected elite core was pre-specified in the "always.selected" constraint of corehunter and selected along with 165 SSD-PGR by the EN algorithm applied to an RD matrix containing both the 200 elite cultivars and the resistant SSD-PGR tail.

The susceptible PGR core was established by minimizing resistant-susceptible SSD-PGR distances and maximizing susceptible SSD-PGR diversity. Bringing resistant and susceptible tails close together should decrease the correlation between phenotypic and genetic similarity, which is in turn expected to increase detection power in association mapping^80^. Therefore, a rectangular $r\times s$ matrix, with $r=$ 165 and $s=$ 249, containing only the RDs between the resistant core and the susceptible tail was generated to identify for each resistant SSD-PGR its closest relative among susceptible SSD-PGR. Here, 86 susceptible SSD-PGR having minimized RD to at least one SSD-PGR from the resistant core were detected (**Supplementary Fig. 5e**). Among them, 55 diverse and less redundant SSD-PGR were selected by applying the EN algorithm to a RD matrix containing only the subset of 86 susceptible SSD-PGR (**Supplementary Fig. 5f**). Without core-selection, the correlation between RD and the absolute differences in $MII$ among 498 resistant plus susceptible SSD-PGR was only 0.05 (Mantel test P-value = 0.011). By naively selecting the 165 most resistant and 55 most susceptible SSD-PGR this correlation would have increased to 0.15 (Mantel test p-value = 0.002). Furthermore, by randomly sampling one hundred times 165 and 55 SSD-PGR from the resistant and susceptible tails, respectively, an average correlation of 0.10 was observed, while correlations were significant in 62% of the cases (Mantel test p-value < 0.05). In contrast, this correlation was 0.00 for the YR-T3C and not significant (Mantel test p-value = 0.44). Afterwards, the selected T3C was adjusted considering SSD-PGR with enough seeds available after selection. The correlation between RD and absolute $MII$ differences virtually did not change due to this adjustment step (*r* = 0.02, Mantel test p-value = 0.34). The final YR-T3C was composed of: 200 diverse European elite cultivars, 150 diverse resistant (${Range}_{MII}$: 0.38%-13.5%, ${Median}_{MII}:$ 6.8%, ${Mean}_{MII}$: 6.9%) and 50 diverse susceptible (${Range}_{MII}$: 27.6%-65.8%, ${Median}_{MII}:$ 36%, ${Mean}_{MII}$: 37.6%) SSD-PGR, with minimized genetic distances among resistant and susceptible PGR pools. Deep sequencing data (for more details see whole genome sequencing in the Methods section) was successfully performed on 191 from the final elite core and 149 YR-resistant and 50 -susceptible from the final SSD-PGR core.

***Extension of the T3C for further traits***

Based on the same strategy deployed for YR, T3Cs were established for 2 further traits, i.e. resistance to LR and powdery mildew (PM) using additional preliminary phenotypic data (not shown). Briefly, each of these two additional cores shares the elite core with the YR-T3C and includes 150 resistant and 50 susceptible SSD-PGR for the respective disease. For these core-sets, however, deep sequencing data for only 28 and 36 diverse SSD-PGR randomly selected from the LR and PM resistant fractions have been generated so far. A list of the 600 SSD-PGR with their respective classifications to the three different T3Cs plus the 200 diverse elite cultivars can be found in **Supplementary Table 15**.

**Additional References (Supplementary Note)**

76.	McIntosh, R. A., Wellings, C. R. & Park, R. F. *Wheat Rusts: An Atlas of Resistance Genes* (CSIRO Publications, 1995).

**Supplementary Figures**

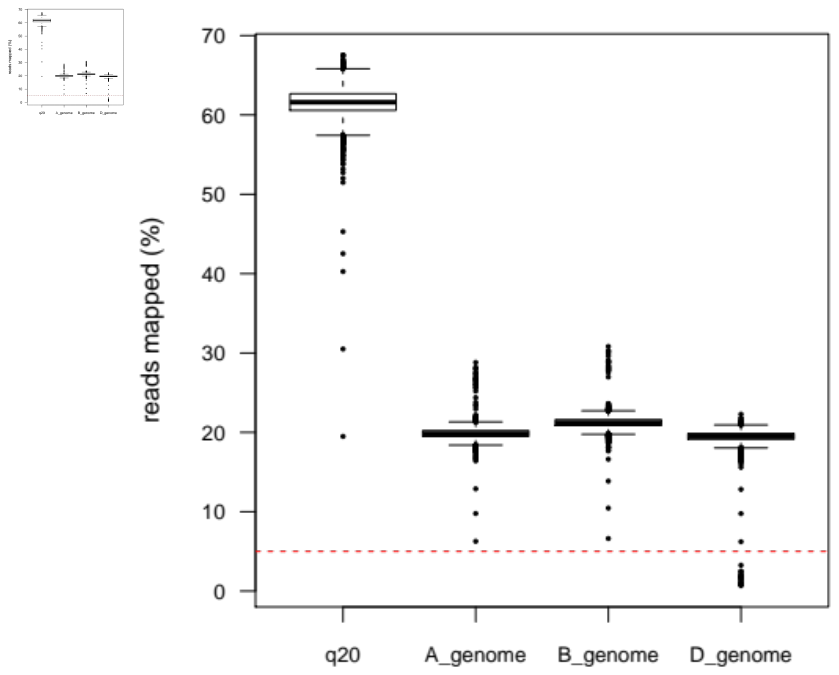

**Supplementary Fig. 1**. Genebank genomics for curating passport data. **a**, Approximately 60% reads were mapped against RefSeq v1.0 with MAPQ 20 wherein ~ 20% reads were mapped on each sub-genome. For 22 accessions, less than 5% reads were mapped on D-genome. These are tetraploid accessions which are potentially falsely labelled as hexaploid (**Supplementary Table 4**). **b**, Normalized read coverage plot for one of the potential tetraploid accessions (TRI 16456). The drop in coverage on D-genome suggest absence of D-genome.

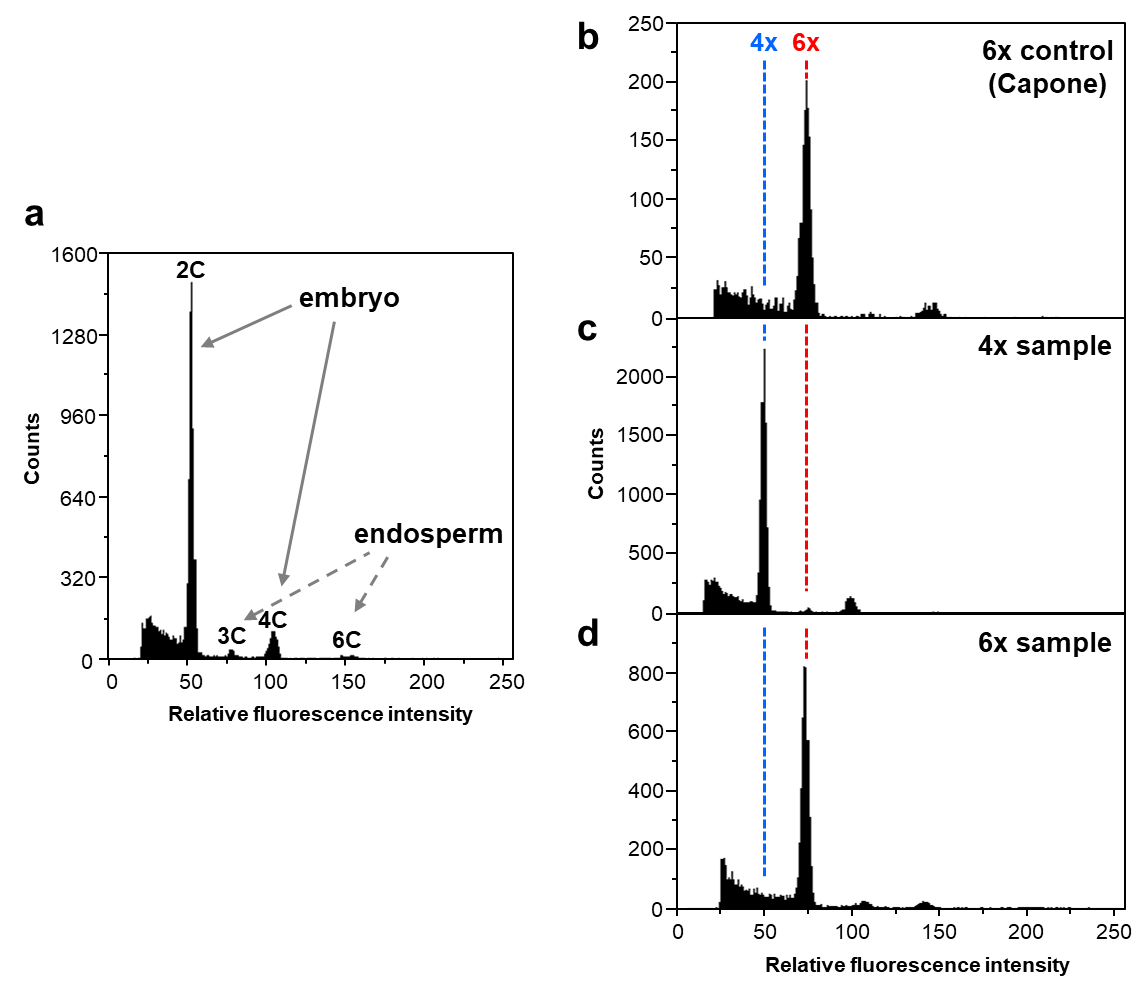

**Supplementary Fig. 2**. Flow cytometric experiments to confirm the tetraploidy (absence of the D genome) for winter wheat plant genetic resources (PGR) from the German Federal ex situ Genebank for Agricultural and Horticultural Crop Species (IPK-genebank) that presented low read coverage for the D genome. **a**, Typical flow cytometric histogram of nuclei isolated from wheat seeds with a pronounced 2C peak (originating from G0/G1 nuclei of the embryo) and a small 3C peak (originating from G0/G1 nuclei of the endosperm). Additional peaks at 4C and 6C represent embryo or endosperm nuclei, respectively, that are either in G2 or endoreduplicated. For the detection of the ploidy level the mean position of the 2C embryo nuclei peaks were compared. **b**, The cultivar Capone (AABBDD) was used as positive control for the presence of the D genome (located at fluorescence channel 75, indicated with a red dashed vertical line) to identify the absence (channel 50, denoted with a blue dashed vertical line) (**c**) or presence (**d**) of the D genome in samples. The complete list of tetraploid accessions falsely labelled as hexaploid wheats can be found in the **Supplementary Table 4**.

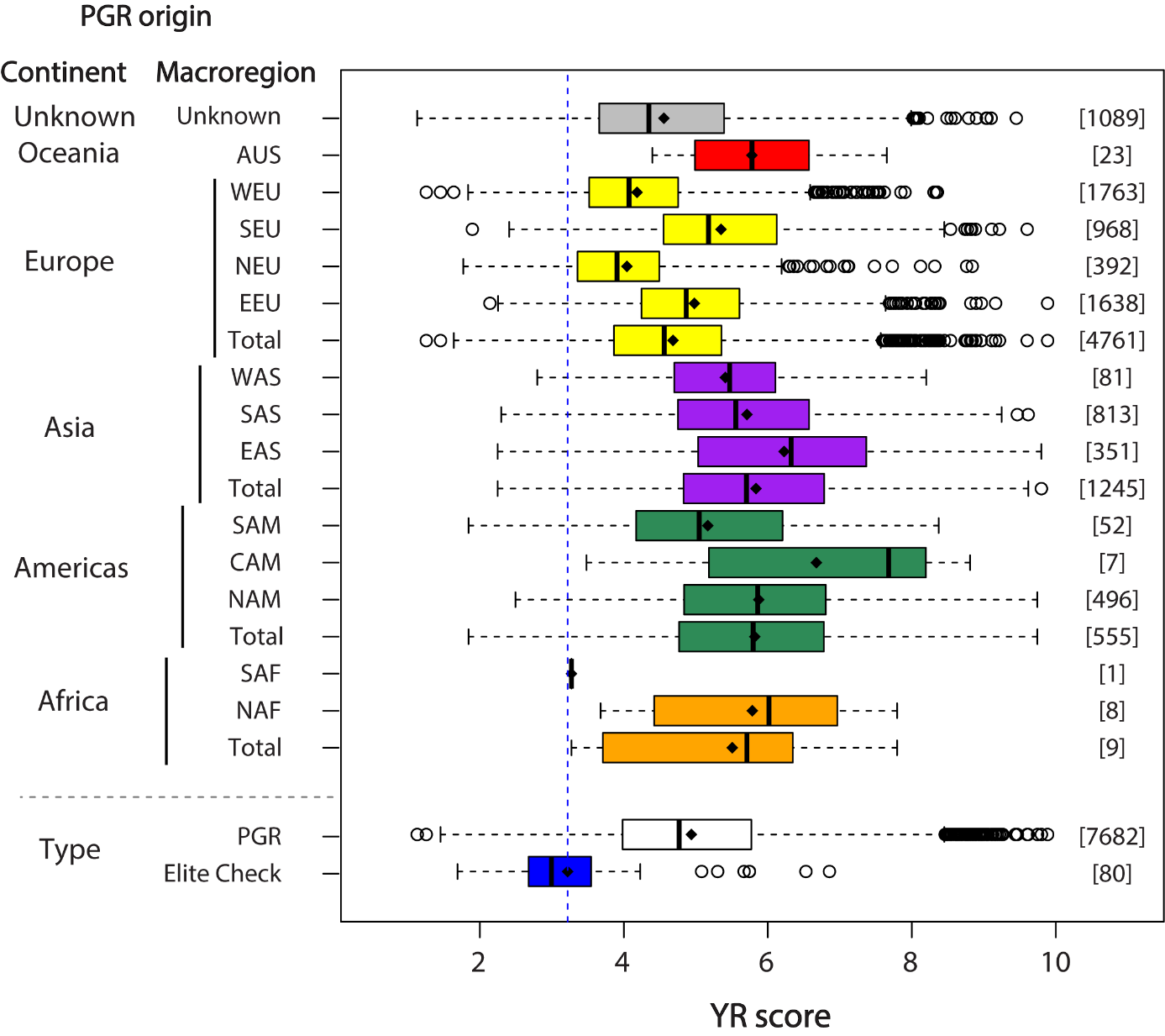

**Supplementary Fig. 3**. Best linear unbiased estimations (BLUEs) for 7,682 plant genetic resources (PGR) of wheat stored at the German Federal ex situ Genebank for Agricultural and Horticultural Crop Species (IPK-genebank) along with 80 European elite cultivar checks screened for resistance against yellow rust (YR, Puccinia striiformis f. sp. tritici) across 12 unbalanced experiments. Experiments were conducted in two German locations between harvest years 2015 and 2020. Resistance was scored based on natural YR infections and using an ordinal rating scale between 1 and 9, where 1 means complete absence of YR leaf symptoms and 9 denotes fully infected leaves. BLUEs that lie outside of the 1-9 parametric space are due to the unorthogonal datastructure of experiments. According to their origins, PGR can be classified into five different continents, namely Africa, Americas, Asia, Europe and Oceania, or their origins are unknown. Continents are further subdivided into 13 megaregions: Northern (NAF) and South (SAF) Africa, Central (CAM), Northern (NAM) and South (SAM) America, Eastern (EAS), Southern (SAS) and Western (WAS) Asia, Eastern (EEU), Northern (NEU), Southern (SEU) and Western (WEU) Europe, as well as Australia and New Zealand (AUS). Black diamonds indicate distribution averages, while [Number] denotes the number of genotypes in each category. The vertical blue dashed line corresponds to the average of check cultivars.

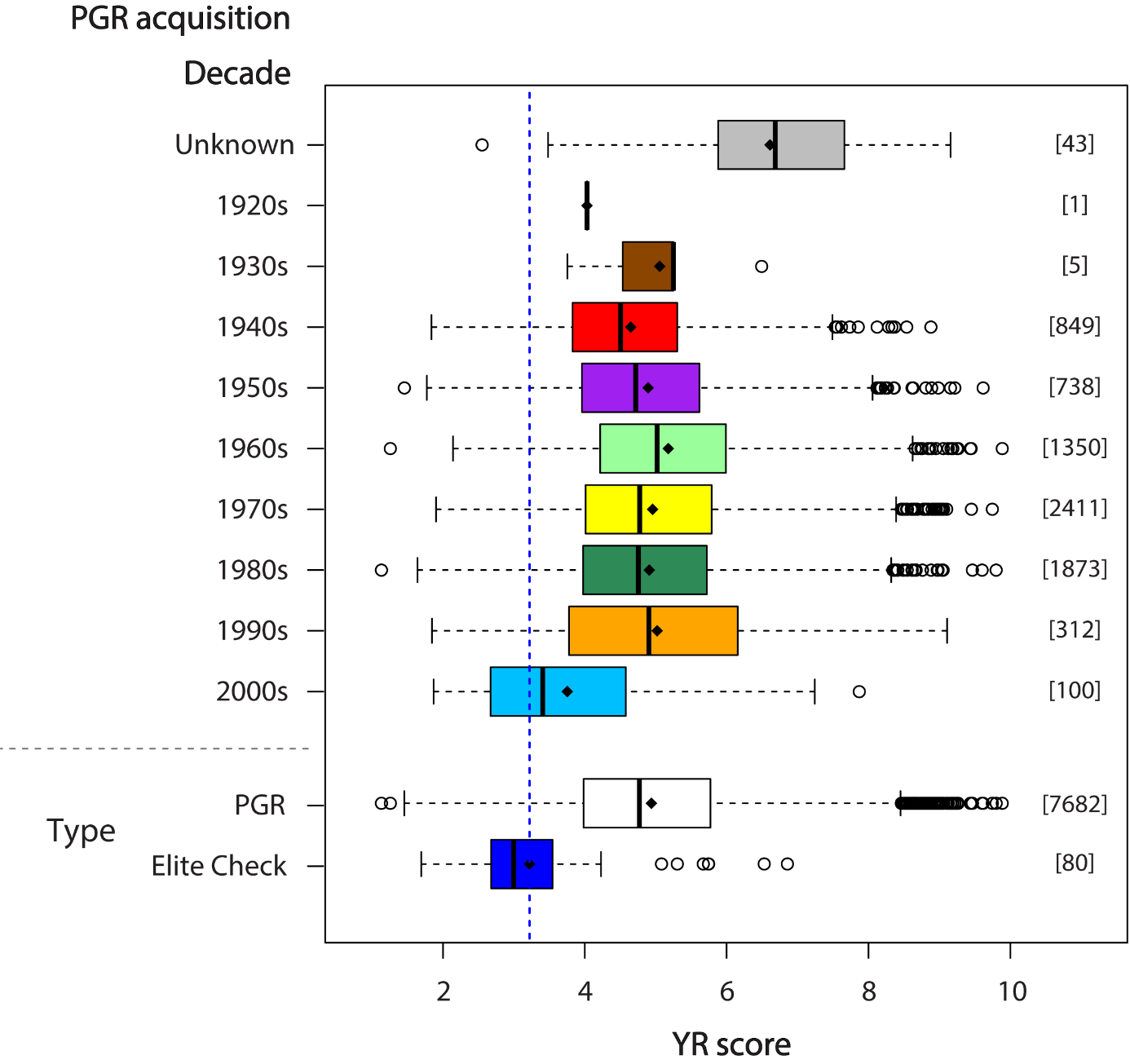

**Supplementary Fig. 4**. Best linear unbiased estimations (BLUEs) for 7,682 plant genetic resources (PGR) of wheat stored at the German Federal ex situ Genebank for Agricultural and Horticultural Crop Species (IPK-genebank) along with 80 European elite cultivar checks screened for resistance against yellow rust (YR, Puccinia striiformis f. sp. tritici) across 12 unbalanced experiments. Experiments were conducted in two German locations between harvest years 2015 and 2020. Resistance was scored based on natural YR infections and using an ordinal rating scale between 1 and 9, where 1 means complete absence of YR leaf symptoms and 9 denotes fully infected leaves. BLUEs that lie outside of the 1-9 parametric space are due to the unorthogonal datastructure of experiments. According to their year of acquisition, PGR can be classified into nine different consecutive decades from the 1920s to the 2000s, or their acquisition years are unknown. Black diamonds indicate distribution averages, while [Number] denotes the number of genotypes in each category. The vertical blue dashed line corresponds to the average of check cultivars.

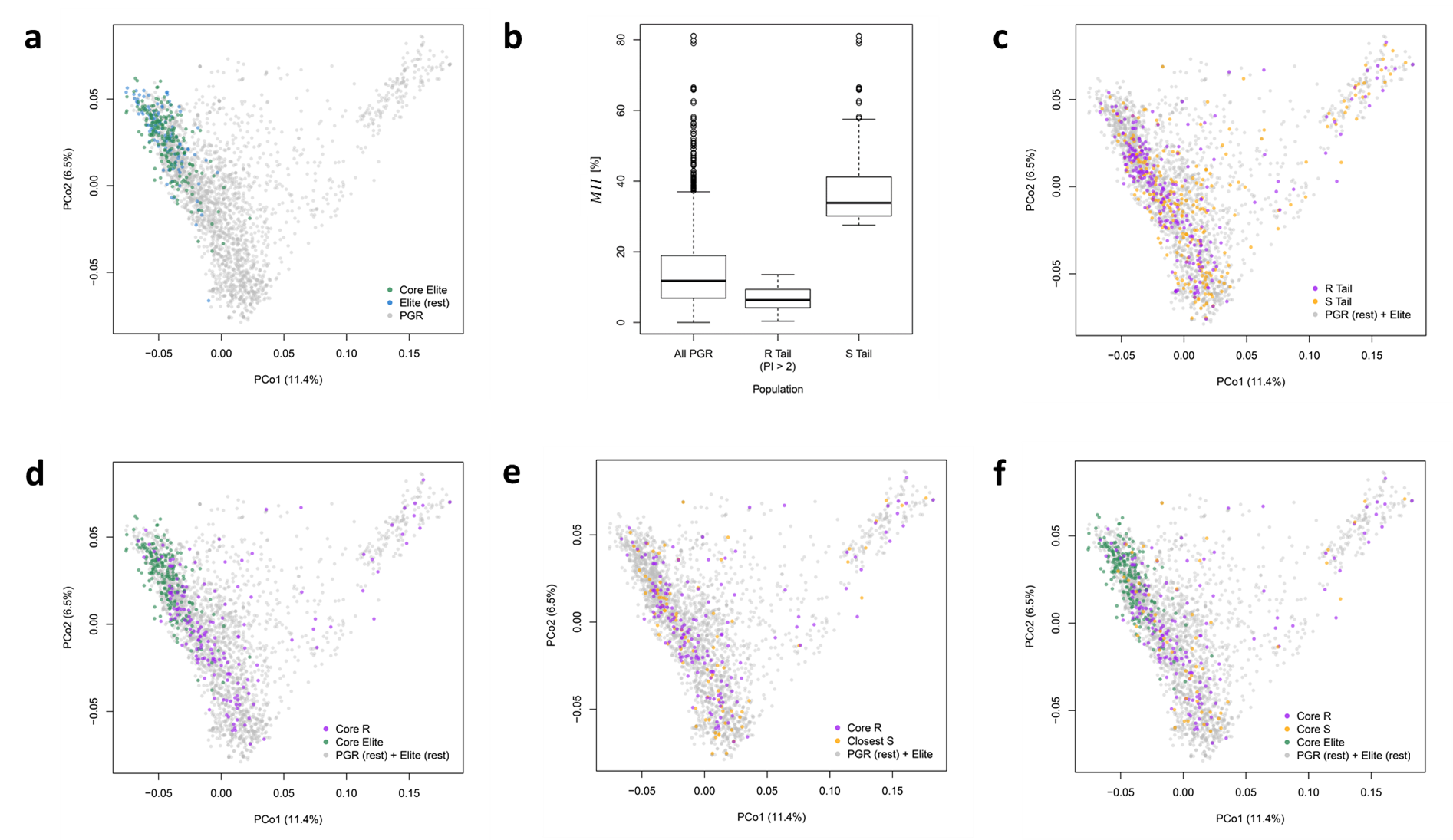
**Supplementary Fig. 5.**

**Supplementary Fig. 5.** Constructing a trait-customized core collection (T3C) to mine for new sources of resistance against yellow rust (YR, *Puccinia striiformis* f. sp. tritici) in wheat accessions (PGR) stored at the German Federal ex situ Genebank for Agricultural and Horticultural Crop Species (IPK-Genebank). **a**, Core-selection of 200 European elite cultivars (green) with maximized diversity and minimized redundancy among a total of 317 cultivars (green and light blue). **b**, Two-tailed selection: the 10% most reliably resistant (phenotyping intensity, PI > 2) and 10% most susceptible PGR were selected among 2,486 PGR isolations (SSD-PGR) according to a YR mean infection index ($MII$ [%]). **c**, SSD-PGR diversity covered by the selected resistant (purple) and susceptible (orange) tails. **d**, Core-selection of 165 SSD-PGR (purple) with maximized diversity and minimized redundancy among SSD-PGR of the resistant tail. **e**, Subset of susceptible SSD-PGR with minimized RD to at least one SSD-PGR from the resistant core that was further reduced to 55 susceptible SSD-PGR with maximized diversity and minimized redundancy. **f**, Selected YR-T3C. All biplots are based on the first two coordinates (PCo1 and 2) from a principal coordinate analysis on the Rogers’ distances calculated among 317 European elite cultivars and 2,486 SSD-PGR. For further details, please refer to the **Supplementary Note**.

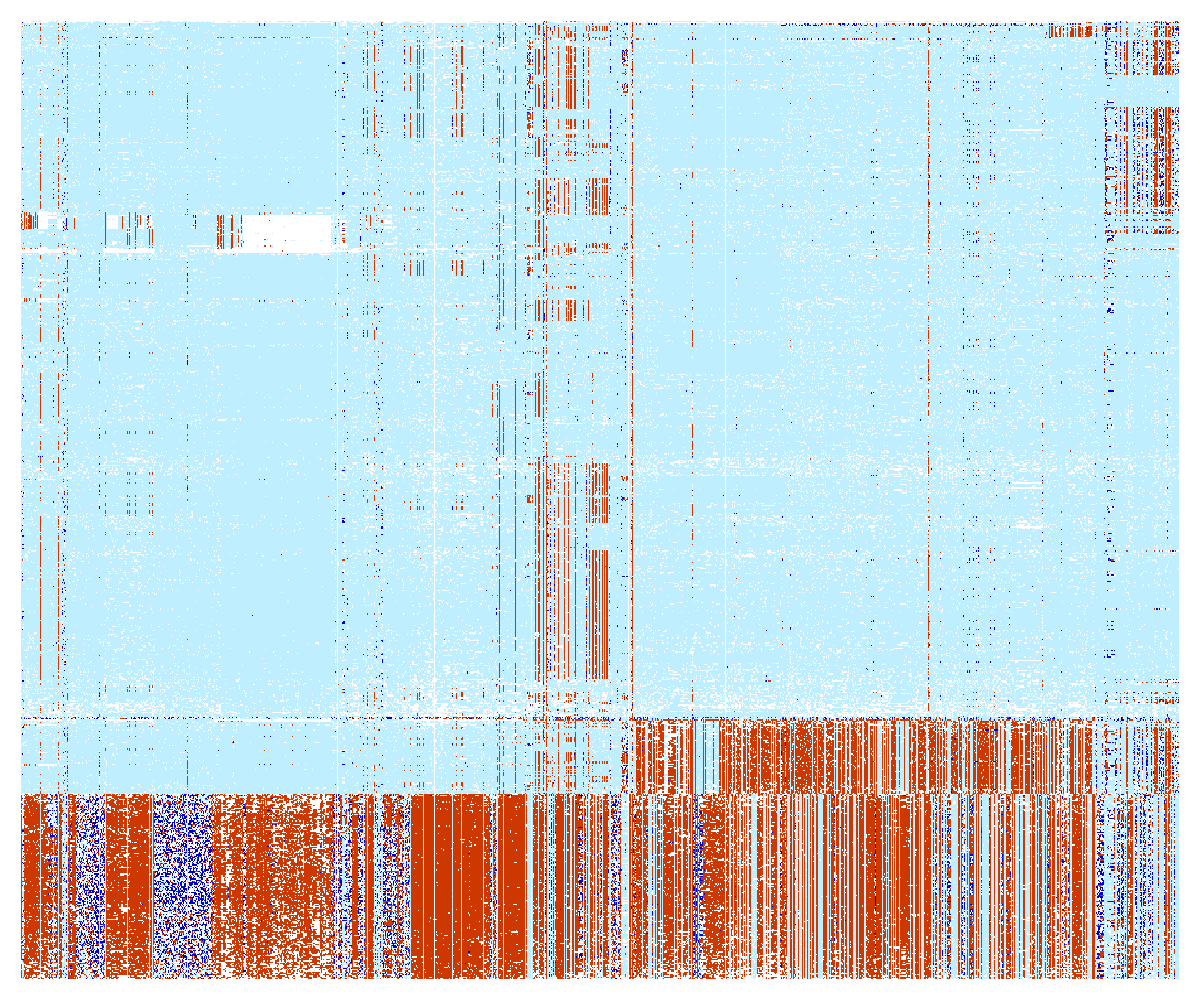

20 Mb

0 Mb

**Supplementary Fig. 6.** Haplotype pattern of 20 Mb region from the short arm of 2A chromosome based on whole genome shotgun sequencing data of the historic panel composed of 760 genotypes: 255 from the Green Revolution (PreGreen), 212 released between 1971 and 2000 (OldCV), and 293 bred after 2000 (NewCV). Each row represents individual genotypes while SNPs are arranged vertically. Alleles identical to RefSeq v1.0 are indicated by blue color; red color represent alternate alleles; heterozygotes are represented by dark blue color while blank spaces represent missing data.

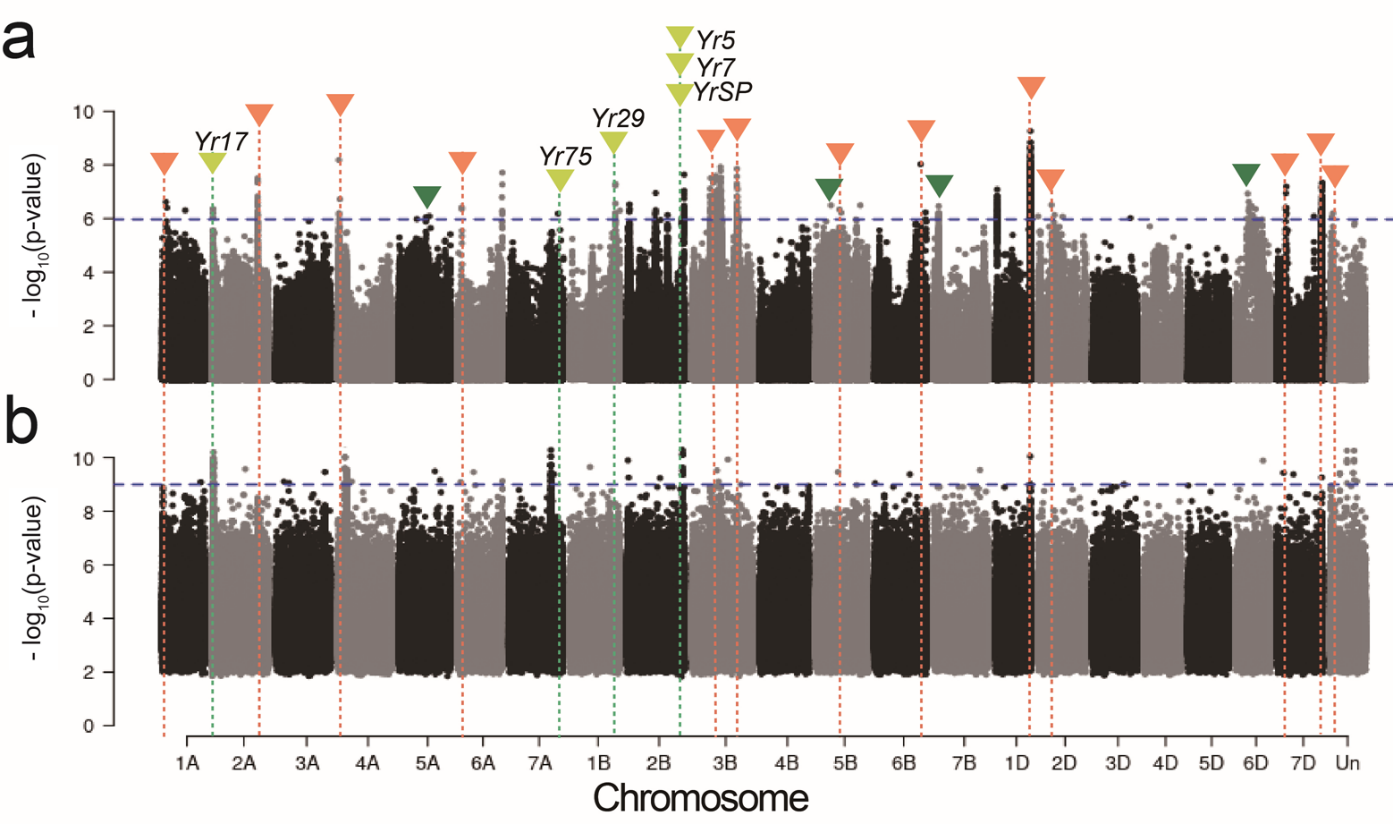

**Supplementary Fig. 7.** Genomic regions harboring loci associated to yellow rust resistance as revealed by (**a**) single-nucleotide-polymorphism-based and (**b**) *k*-mer-based GWAS in the whole panel (elite cultivars + T3C PGR). The -log_10_(p-value)s are shown on the y-axis while all whole genome shotgun reads portrayed in Manhattan plots were mapped back to Chinese Spring v1.0. Significance thresholds for associations are denoted with blue horizontal dashed lines. The positions of known YR resistance loci are indicated with lime vertical dashed lines, while associated regions compromising loci where most resistance-conferring alleles are contributed by PGR are shown using an orange dashed line. The green triangles mean the resistance-conferring alleles are almost fixed in elite cultivars.

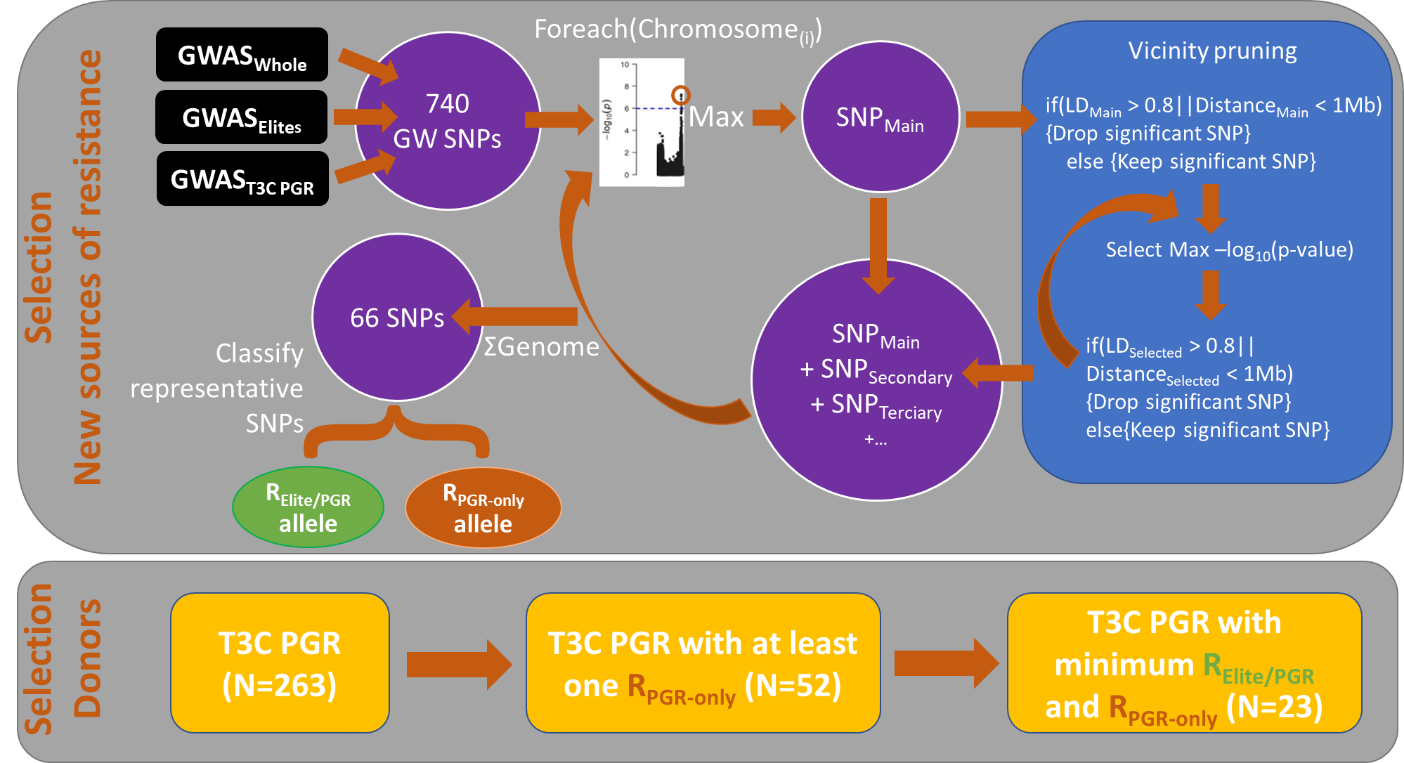

**Supplementary Fig. 8.** Workflow of resistance donor selection. The 740 significant SNPs resulting from genome-wide association (GWAS) scans for yellow rust resistance (**Supplementary Table 22**) were reduced to 66 representative SNPs by considering most significant peaks per chromosome and pruning significant SNPs in their vicinity with high linkage disequilibrium (*r*^2^ > 0.8) or mapping proximal to them (< 1 Mb). The 66 representative significant SNPs (**Supplementary Table 24**) were classified according to their presence - R_Elite/PGR_ - or complete absence - R_PGR-only_ - in the elite genepool. Among the 263 possible PGR from the trait customized core collections (T3C), 52 contain at least one R_PGR-only_ allele at a representative SNP (**Supplementary Table 28**). The set of 52 donors can be further reduced to 23 T3C PGR with minimized R_Elite/PGR_ and R_PGR-only,_ which correspond to the best near-isogenic line proxies found for 30 new sources of YR resistance (**Supplementary Table 27**).

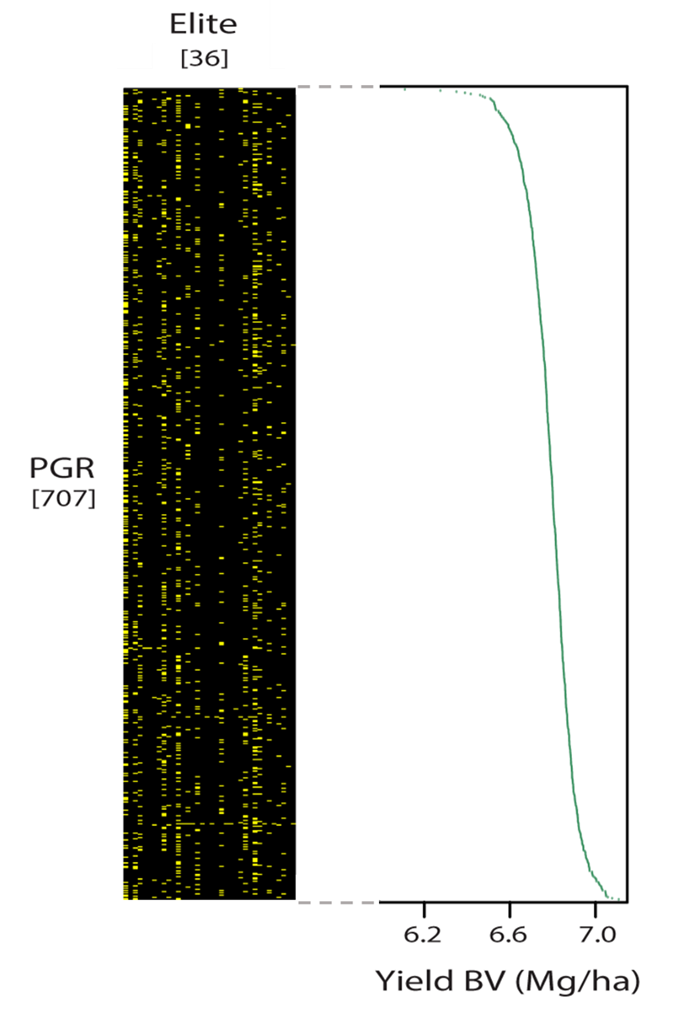

**Supplementary Fig. 9.** Estimation of the yield breeding value for plant genetic resources (PGR) using an Elite×PGR hybrid concept. An incomplete factorial mating design between 707 PGR and 36 elite cultivars allowed the obtention of 1,427 Elite×PGR F_1_ hybrids (left, indicated in yellow). In crossing blocks, PGR served as male pollen donors, while female elite cultivars were obtained using gametocide applications during hybrid seed production. Yield performances of hybrids across multiple-environments were used afterwards to estimate the breeding value (Mg/ha) of each PGR parent (right). The distribution of yield breeding values is indicated with green dots.

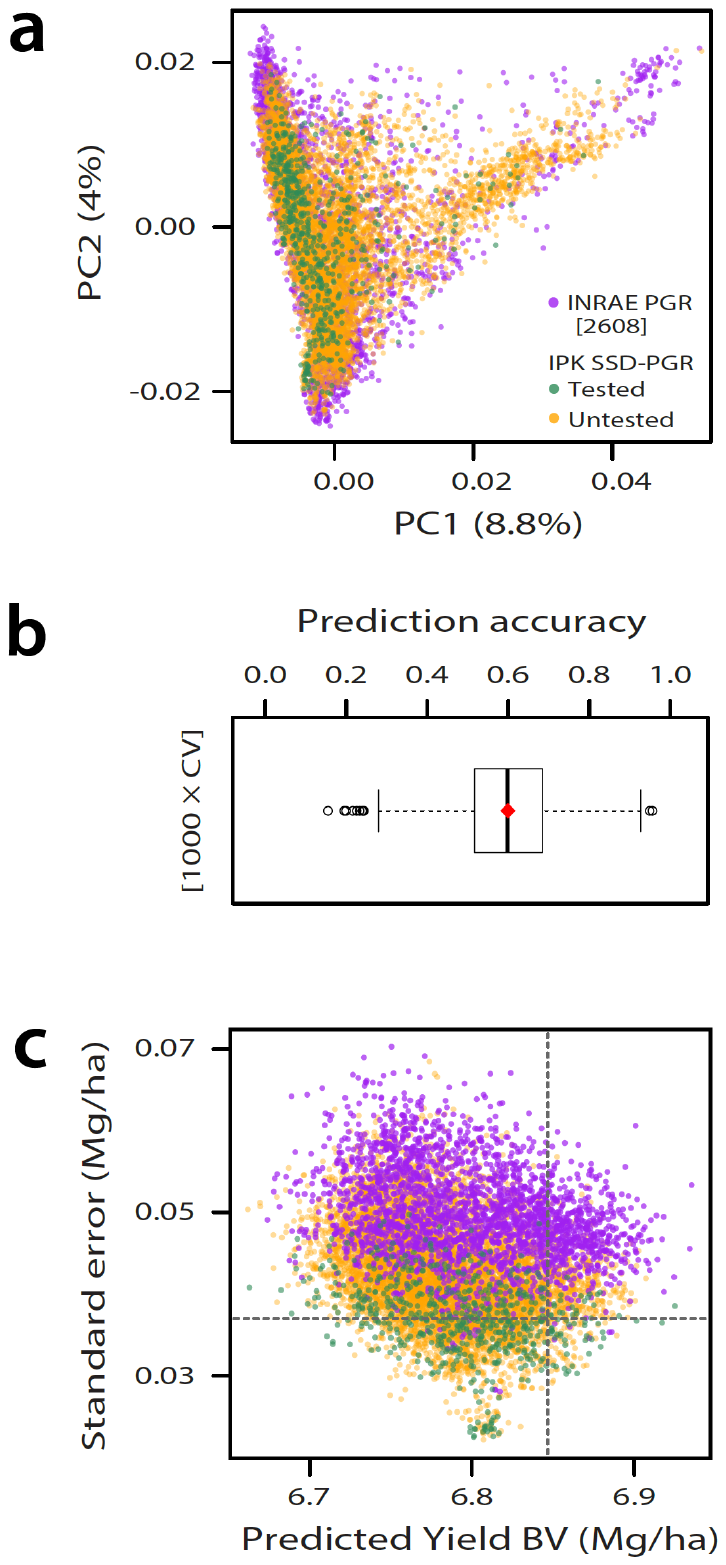

**Supplementary Fig. 10.** Genomic prediction of yield breeding values (BV, Mg/ha) of plant genetic resources (PGR) across IPK and INRAE genebanks. **a**, Neutral molecular diversity portrayed by the first two principal components from the merged SNP matrix (895 common variants). Five hundred ninety-seven PGR isolations (SSD-PGR) have yield BV estimates (tested, in green) and conformed the training set for genomic prediction of 7,148 untested SSD-PGR (in orange) from IPK and 2,608 PGR from the INRAE genebank (purple). BV were estimated using an Elite$\times$PGR hybrid concept (see **Supplementary Fig. 9**). **b**, Yield BV prediction accuracy of GBLUP in 1,000 five-fold cross-validation runs within the training set using the 895 common SNP variants between genebanks. The red diamond indicates the average. **c**, Yield BV genomic predictions and their standard errors (SE) for IPK genebank and INRAE genebanks. The vertical dashed line indicates the 10% superior yielding predictions while the horizontal one is the average SE of the training set.
